# The Influenza Hemagglutinin Cytoplasmic Tail Domain Interacts with Phosphatidylinositol 4,5-bisphosphate

**DOI:** 10.64898/2026.08.24.746733

**Authors:** David Winski, Matthew T. Parent, Jaqulin N. Wallace, Chamithri Weerakoon, Suman Shrestha, Prakash Raut, Hang Waters, Joshua Zimmerberg, Alexander Sodt, Samuel T. Hess

**Affiliations:** Department of Physics and Astronomy, 120 Bennett Hall, University of Maine, Orono, ME 04469-5709, USA; Section on Integrative Biophysics, Eunice Kennedy Shriver National Institute of Child Health and Human Development, National Institutes of Health, 10 Center Drive, Room 10D14, MSC 1855, Bethesda MD 20892- 1855, USA; Section on Membrane Chemical Physics, Eunice Kennedy Shriver National Institute of Child Health and Human Development, National Institutes of Health, 9000 Rockville Pike, Bethesda MD 20892 USA

## Abstract

During the influenza viral life cycle, the viral glycoprotein hemagglutinin (HA) mediates binding, entry, and fusion. Densely packed clusters of HA trimers at the plasma membrane are required to produce infectious virions; however, the mechanism of HA clustering is still unknown. We have shown previously that HA co-clusters with and modulates phosphatidylinositol 4,5-bisphosphate (PIP2) in host cell plasma membranes (PM). Here, we further characterize the relationship between HA and PIP2 using molecular dynamics simulations (MD) and fluorescence photoactivation localization microscopy (FPALM) to elucidate a mechanism of HA-PIP2 interaction. We found that the interaction occurs largely between the PIP2 head group and the cytoplasmic tail domain (CTD) of HA. Mutations of the CTD were made to alter charge (HARE, HARREQ), palmitoylation sites (HAMAY), or a combination thereof (HAREMAY, RREQMAY). MD showed that HARREQ and RREQMAY had the strongest effect on HA-PIP2 interactions through a depletion in the radial distribution function of PIP2 around HA at distances ≤2.5 nm. FPALM revealed that HA cluster density at the PM was significantly reduced by CTD mutations, with the largest reduction occurring in mutants where the CTD charge and acylation were both altered (HAREMAY). HAREMAY clusters were also found to have larger circularities and perimeters, implying a structural change to the clusters. Mutations in the HA transmembrane domain also caused modest changes to the cluster properties of HA and its co-clustering with PIP2. FPALM showed PIP2 co-clustering with HA was also affected by HA mutations with more free PIP2 localized under HAREMAY clusters. A chemical model of simultaneous HA-PIP2 and PH-PIP2 binding enables interpretation of HA-PIP2 interactions and reveals quantitative differences between PIP2 binding by HA CTD mutants. We conclude that the mechanism of HA-PIP2 interaction consists of at least electrostatic and hydrophobic components. Our insights into the mechanism of HA-PIP2 interaction, and the prevalence of putative PIP2-interacting domains in a number of viral spike proteins suggest it may be fruitful to identify methods of disrupting interactions between phosphoinositides and viral proteins.

## INTRODUCTION

Influenza virus (IAV) remains a public health threat which causes thousands of deaths in the United States each year [1]. Due to its high mutation rate, a universal treatment option has not yet been created. IAV infects humans by exploiting host cell membrane organization [2]. An improved understanding of the processes underlying membrane organization is needed.

IAV forms enveloped viral particles consisting of a lipid bilayer derived from the host cell and enriched in certain lipids [3–6]. Anchored in the IAV lipid envelope are the two viral glycoproteins, hemagglutinin (HA) and neuraminidase (NA), and the matrix-2 (M2) ion channel protein [7, 8]. IAV entry into host cells depends on membrane fusion, catalyzed by HA [2]. Clustering of IAV membrane proteins also mediates release of vRNA from the nucleus [9] and recruits M1 for multimerization beneath the PM [10]. HA-dependent membrane fusion relies on high density of HA trimer clustering [4, 11]; however, the mechanism for HA clustering is unknown.

HA is composed of three subunits: the ectodomain, transmembrane domain (TMD), and cytoplasmic tail domain (CTD) [7]. While the TMD and ectodomain are relatively large, the CTD is only 10-11 amino acids long, extending into the cytoplasm close to the plasma membrane (PM), and is presumably anchored to the membrane by three cysteines that are typically acylated[12] with either a palmitic or stearic acid[13]. The tail has been proposed to mediate assembly of other viral components, specifically M1 and NA[14–16]. Mutation of the CTD cysteines disrupts HA association with M1[6, 17], and inhibits viral growth,[6, 18]; alternately, they revert back to a cysteine during live infection[17, 19]. Mutations of the TMD and CTD decreased HA clustering at the PM, which in turn decreased infection rates[4, 19–21]. Thus, we suspected the CTD could play a role in HA clustering.

HA CTD palmitoylation is essential for influenza infectivity[6, 17, 22]. Palmitoylation of the HA CTD alters membrane curvature in virus-like particles[17], which may reduce the energy barrier during the formation of viral buds. Because of the high level of conservation of the HA CTD cysteines and their importance to infectivity[23], these make promising targets for a “universal therapy”[24].

Furthermore, the CTD of HA consistently contains amino acid sequence motifs with at least 1 positive residue adjacent to one of the acylated cysteines[25]. We hypothesized that this motif could interact with phosphatidylinositol 4,5-bisphosphate (PIP2), which has been previously observed to be co-localized with HA[26]. In general, PIP2 comprises only about 1% of the total lipid composition of the PM[27] but serves a diverse range of important functions for the cell, especially in signaling, regulation of membrane trafficking, and control of the cytoskeleton[28]. Considering that HA clusters are frequently observed together with actin rich membrane regions of the PM[29, 30], that actin and actin binding proteins (ABPs) are found in virions[31], that most of those ABPs also have known PIP2 association[32, 33], and that diffusion of a fluorescent PIP2 analog is modulated by the presence of HA[26], PIP2 may serve as a mediator between influenza and the actin cytoskeleton.

Other viruses are known to exploit PIP2 to form clusters on the PM of infected cells. Examples include HIV gag[34–36] and Ebola VP40[36–38] structural proteins, which both interact with PIP2 at the PM. Platforms of gag on the PM are completely lost upon depletion of PIP2, and PIP2 is required for successful HIV assembly[39]. HIV gag proteins have been shown to directly interact with PIP2 through a variety of basic residues present in the matrix domain, nearest the inner leaflet of the PM[40, 41]. In the case of the Ebola virus, VP40 is thought to interact with PIP2 through cationic side chains of the VP40 protein in order to stabilize clusters[37]. Influenza may follow a similar pattern to allow effective clustering of its proteins on the PM for assembly prior to viral budding.

We hypothesize that the cytoplasmic tail domain (CTD) of HA can interact with phosphoinositides such as PIP2. To test the roles of CTD palmitoylation and charge on HA clustering at the PM and co-clustering with PIP2 (PH domain), we 1) used molecular dynamics (MD) simulations of HA in bilayers containing PIP2 and 2) used fluorescence photoactivation localization microscopy (FPALM) to image Dendra2-HA and PAmKate-PH(PLCδ) for wild-type HA (HAwt) and for HA mutants that removed CTD palmitoylation sites (HAMAY), altered CTD charge (HARE and HARREQ), or both (HAREMAY). HA mutants are summarized in Table 1. Our findings help illuminate a mechanism for HA-PIP2 interaction.

**Table 1.**
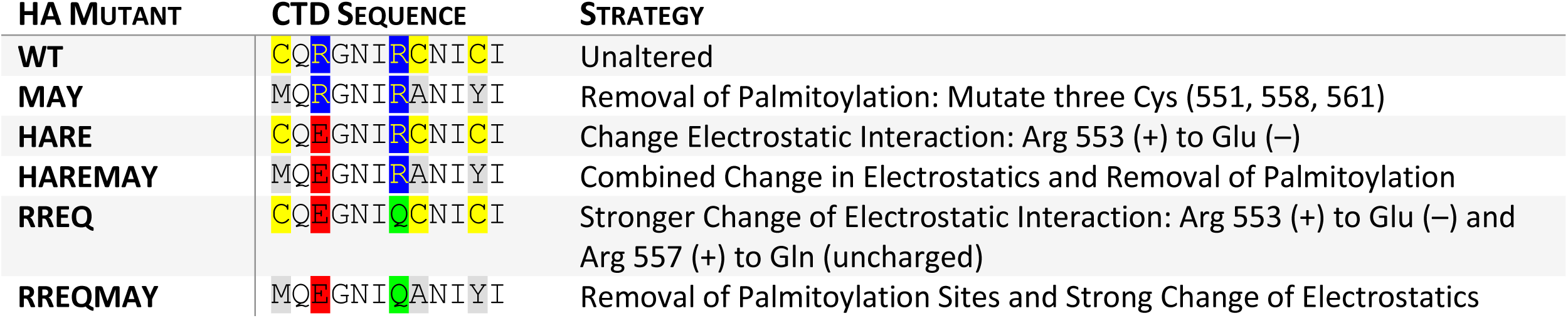
List of HA CTD mutations that were used in the MD simulations and the FPALM experiments. Each mutant is listed with the amino acid sequence and the effect the mutation has on the CTD.

## MATERIALS AND METHODS

### Cell Culture

NIH3T3 mouse fibroblast cells (ATCC, CRL-1658) were cultured in T25 Nunc™ flasks (Thermo Scientific, 136196) in Dulbecco’s Modified Eagle Medium (DMEM) (Lonza, 12-604F) supplemented with 10% calf bovine serum (ATCC, 30-2030), 100 µg/mL penicillin-streptomycin, and incubated at 37°C and 5% CO_2_. Cells were not allowed to reach more than 80% confluency before being passaged to a new flask.

### Plasmids

Expression vectors for HA (X-31B, Puerto Rico/8/1934-Aichi/2/1968) were constructed to fuse the HA with Dendra2 at the N-terminal of the HA (Dendra2-HA) using previously published methods[26, 42]. Mutant HAs were created to alter the charge and/or remove palmitoylation sites within the cytoplasmic domain (CTD). To eliminate CTD palmitoylation, mutations of Cys 555, Cys 562, and Cys 565 were made into Met 555, Ala 562, and Tyr 565, producing a mutant called HAMAY using a QuikChange II Site-Directed Mutagenesis kit (Agilent, cat #200523) as previously published[17]. Dendra2-HAMAY was created by cloning Dendra2 into the N terminal of the mutated HAMAY using basic cloning techniques similar to published methods used for the creation of Dendra2-HA[26, 42]. In a similar method used for the creation of the mutant HAMAY, the following mutations were done for the remaining HA fusion proteins: to produce a neutrally charged CTD, Arg 561 was mutated by site directed mutagenesis into Glu 561 in the fusion protein, Dendra2-HA, to produce the mutant variant, Dendra2-HARE; to have a CTD with no palmitoylations and be neutrally charged Arg 561 was mutated by site directed mutagenesis into Glu 561 in the fusion protein, Dendra2-HAMAY, to produce the mutant variant, Dendra2-HAREMAY; and to produce a negatively charged CTD, Arg 557 and Arg 561 were mutated by site directed mutagenesis into Glu 557 and Gln 561 in the fusion protein, Dendra2-HA, to produce the mutant variant, Dendra2-HARREQ.

As shown previously[26], to visualize PIP2 in the PM we used the pleckstrin-homology (PH) domain from PLCä[43-46] fused to PAmKate[47].

### Transient Transfection and Fixation

For imaging, cells were plated onto 35mm petri dishes with a #1.5 coverglass bottom (MatTek, P35G-1.5-20-C) in complete growth medium (without antibiotics and without phenol red) and seeded at a concentration of 7-8×10^4^ cells/plate. After 24 hours of growth, cells were transfected using Lipofectamine 3000 (Invitrogen, L3000008) with a total of 2 µg of plasmid DNA, 4 µL of P3000, and 7.5 µL of Lipofectamine per plate per dish. Transfected cells were covered with aluminum foil to prevent activation of the fluorescent proteins within the incubator or by room lights and left to grow for another 24-36 hours in the incubator. Cells were then washed with phosphate buffered saline (PBS) (Sigma-Aldrich, D8537) three times to remove growth media and detached cells, fixed at room temperature in 4% paraformaldehyde (PFA) (Alfa Aesar, J61899AK) for 10 minutes, and then washed again with PBS three times.

### Super-Resolution Microscopy

Two-color FPALM was performed as described previously[47]. Briefly, a 558 nm readout (CrystaLaser, 100 mW), and a 405 nm activation (CrystaLaser, 5 mW), were combined using a dichroic mirror (Chroma, Z405RDC) and are focused through a lens (ThorLabs, f=350 mm) to reach a focus in the back aperture of the objective lens (Olympus 60x, 1.45 NA oil) mounted on an inverted microscope (Olympus, IX71). The 558 nm laser was operated with peak intensity 1.0 kW/cm^2^ ± 0.1 kW/cm^2^ at the center of the field, and the 405 nm laser was operated at low (but variable intensity), typically less than 0.02 W/cm^2^, as measured at the sample. For imaging of PA-JF646 samples, a 638 nm laser (CrystaLaser, 100mW) was added and aligned to the 558 nm and 405 nm lasers and operated at 0.8 kW/cm^2^ ± 0.1 kW/cm^2^ peak intensity at the sample. The activation laser passes through a half wave plate (Newport, 10RP42-1) and a linear polarizer (Newport, 5511) is used to match the polarization of the activation laser to the readout laser. Then, both lasers are passed through a quarter wave plate (Newport, 10RP54-1B) to obtain elliptical (approximately circular) polarization of the light at the sample. A quad band dichroic mirror DM1 (Semrock, Di01-R405/488/561/635) in the turret of the microscope reflects the laser light into the objective, which changes the wavefront curvature to produce nearly parallel illumination of the sample (widefield) or with translation of the illumination beam off axis to produce total internal reflection at the coverslip-sample interface for excitation of fluorescence (TIRF) just above the coverslip.

Fluorescence collected by the objective passes through DM1, through the tube lens, and into a 2x telescope (f=400 mm, f=200 mm). After the telescope, the fluorescence reaches an ultra-flat dichroic (Semrock, FF580-FDi02-t3), splitting the light into a reflected and transmitted channel which are filtered (Transmitted channel: Chroma, ET605/70m; Reflected channel: Semrock, FF01-585/40-25) before forming adjacent images on the sensor of an EMCCD (Andor, iXon 3). Acquisitions were typically at least 10,000 frames at 31 Hz with an EM gain of 200. Using a mercury arc lamp (Olympus, U-RFL-T) filtered with an excitation filter (Chroma 476/10X), cells were selected for moderate expression of Dendra2 (via green fluorescence).

### Localization, Bleed-Through Correction, and Drift Correction

Localization was performed by MATLAB scripts which 1. background subtract using a rolling ball (6 pixel radius)[48] or temporal median algorithm (100 frame window)[49], 2. identify individual molecules by threshold, 3. fit molecular images with a Gaussian to determine x and y positions, Gaussian 1/e^2^ radius, and amplitude. *Fit Quality Control (“Tolerances.”)*: Fits which fell far beyond the normal range of values for Gaussian radius, localization precision, number of photons detected, or fractional errors in the radius and Gaussian amplitude were deemed to not pass tolerances and not used for subsequent analysis. The 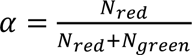 value was determined for each localized molecule to determine the molecular species[47], where N_red_ and N_green_ are the number of detected photons in the “red” (long-wavelength) and “green” (short-wavelength) detection channels, respectively. Bleed-through correction was applied as described previously[50]; the ranges of *α* used to define each species was chosen such that the bleed-through rate was <10% for both Dendra2 and PAmKate. Molecular coordinates were drift corrected following the method of Mlodzianoski et al. [51] binning molecules onto a grid (spacing 100 nm) in temporal groups “chunks” each containing ∼10,000 molecules. Chunks are then translated to optimize cross correlation of successive chunks. Translations in x and y resulting in best cross-correlation were used to compute drift using a 2nd order polynomial which could be applied directly to localized coordinates on a per frame basis[51].

### Combination of Successive Localizations (CSL)

Localizations which fell within successive frames and were within ±f_xy_ (the localization precision) of one another were combined into a single localization with position equal to the intensity-weighted average of the two (or more) localizations, and the number of photons equal to the sum of photons from all combined data.

### Image Rendering

Localized molecules for each species were plotted using intensity-weighted gaussians of prescribed size (*σ* = 20*nm*) for all molecules. Hemagglutinin-Dendra2 molecules were rendered in green and PH domain-PAmKate molecules were rendered in magenta. Overlap in molecules, signifying colocalization, was represented in grey or white. Unless otherwise noted, scale bars are 1 μm.

### Data Analysis for Clustering and Colocalization

Localized coordinates for individual molecules were analyzed to determine radial distribution functions, and clustering properties. Analysis of localization, cluster identification, trajectory analysis, CSL, and radial distribution function (RDF)[52] calculation was performed using custom scripts written in Matlab and executed in version 2016b or later. The RDF of molecule A, i.e. g_A_(r), quantifies the probability of observing molecule A as a function of distance r from an average molecule of A. RDF is normalized such that the amplitude is 1 when the probability of A is equal to the probability expected for a uniform distribution; amplitudes larger than 1 indicate enrichment, and amplitudes less than 1 indicate depletion. RDFs for two different molecular species, e.g. g_AB_(r) calculate the probability of finding molecule B as a function of distance from an average molecule A, with the same normalization that g_AB_(r) = 1 when the probability equals that expected for a uniform distribution. Cluster-based RDFs, i.e. g_AB_(r), calculate the probability of observing a molecule of B as a function of distance r from the center of a *cluster* of A[26]. Measurements of camera pixel size and confocal figures were performed in Fiji, a distribution of ImageJ.

### Density-Based Clustering

Clusters were identified from localized molecular coordinates by previously published methods[26, 29]. Briefly, localization data for each of two species were split by α values (see FPALM methods) and binned into grids with (10nm)^2^ pixels, yielding one grid for each species (red or green). Each grid was then convolved with a uniform circular disk of radius 50nm. A mask was then automatically calculated (see supplemental methods) and the cell-averaged density (number of localizations per unit area) of each species determined. To identify clusters, the convolved green or red (Dendra2 or PAmKate, respectively) localization grid was thresholded for all contiguous regions above either 3 or 4 times the average density for that cell. Contiguous pixel areas were then quantified using the MATLAB regionprops function to determine cluster area and density of both species. Density values determined using this method were reported as “relative to average density” in this text. Low and high concentrations of a given species within clusters of the other were defined as “low” when below the average for the given species, and “high” when ≥3 times the average for that species.

### Significance Testing

Significance testing was performed using GraphPad Prism 8.3.1 software using data imported from Matlab or other analysis. Significance p-value cut offs were indicated as follows, with increasing significance, p≥0.05 (ns = not significant), p<0.05 (*), p<0.01 (**), p<0.001 (***), and p<0.0001 (****). The p-value specifies the probability that the test cannot reject the null hypothesis, that the two groups come from the same underlying distribution. Exact p-values can be found in the appendix. All nonlinear fits (other than localization) were done using Microcal™ Origin 6.0 software. Unless otherwise noted, error bars denote the 95% confidence interval and are computed using the standard deviation of the experimental data.

### Molecular Dynamics Simulations

All simulations were run in the CHARMM36m forcefield using the AMBER[53] program package. The structure of HA from RCSB 6HJR[54] was used, which consists of the ectodomain and TMD. The secondary structure of the CTD is not well defined. Based on AlphaFold[55] predictions, the tail was modeled as an alpha helix using the known sequence of the CTD for IAV. For all HA simulations, the ectodomain was also removed to decrease the number of molecules in the system to improve efficiency as preliminary data was taken to show that removing the ectodomain did not affect the interaction of HA with PIP2. The removal of the ectodomain along with modeling the CTD was done in PyMOL[56]. The three cystines found in the final 12 amino acids of the c-terminal of HA were palmitoylated to mimic physiological HA. The protein was embedded in a lipid bilayer that mimics physiological cellular plasma membrane using CHARMM-GUI[57–61]. The extracellular leaflet consisted of 35% cholesterol, 31% 1,2-Dioleoyl-sn-glycero-3-phosphocholine (DOPC), 29% Sphingomyelin (DSM), 3% 1,2-Dioleoyl-sn-glycero-3- phosphoethanolamine (DOPE). The cytoplasmic leaflet consisted of 38% cholesterol, 17% DOPC, 16% DSM, 15% DOPE, 3% 1,2-dioleoyl-sn-glycero-3-phospho-L-serine (DOPS), and 10% PIP2. Na^+^ and Cl^-^ ions were added to neutralize the system with an effective final NaCl concentration of 150mM. Water was modeled with TIP3, as is appropriate for the CHARMM forcefield. HA mutant systems were created to copy the sequence mutations found in experimental data.

## RESULTS

We hypothesize that HA can interact with PIP2 through its cytoplasmic tail domain (CTD). We therefore investigated how changes in the CTD affected HA clustering and HA co-clustering with PIP2. To do this, we designed mutations to the CTD that alter its charge and/or remove the palmitoylation sites (Table 1). To test whether the palmitoylation plays a role in the interaction with PIP2, the three cystines known to be palmitoylated were mutated to methionine (M), alanine (A), and tyrosine (Y), which has been previously named HAMAY[17]. To test for electrostatic interaction, two mutants were constructed: HARE mutates the first arginine (positively charged) to glutamic acid (negatively charged, E), which presumably gives the CTD a net charge of zero. HARREQ takes HARE and adds a mutation of the second arginine to glutamine (neutral, Q) which presumably gives the CTD a net negative charge. HAREMAY and HARREQMAY were constructed to combine MAY with HARE or HARREQ, respectively, testing the combination of both electrostatic and acylation changes to the CTD.

### Molecular dynamics simulations of HA in solvated bilayers elucidate an HA-PIP2 interaction

To understand how each CTD mutation affects the HA-PIP2 interaction, all-atom molecular dynamics (MD) simulations were run for each of the HA mutants in Table 1. A single HA trimer was placed in a lipid bilayer approximating a mammalian PM in composition. The amount of PIP2 in the cytoplasmic leaflet was increased so that the rate of any interactions between HA and PIP2 was high enough to sample those interactions within the timescale simulated. HAwt can be seen to directly interact with PIP2 (Figure 1). Each arginine on the CTD is within 0.5 nm ± 0.02 nm of phosphates 4 and 5 on the PIP2 headgroup. The PIP2 headgroup interacts with the HA CTD for times comparable to the simulation length itself: off rate analysis was characterized using Pylipid, resulting in a large range (0.001 to 0.106 ns^-1^ for Arg 213, and 0.001-0.473 ns^-1^ for Arg 217), suggesting residence times of a few to hundreds of ns for PIP2 bound to HA.

**Figure 1.**
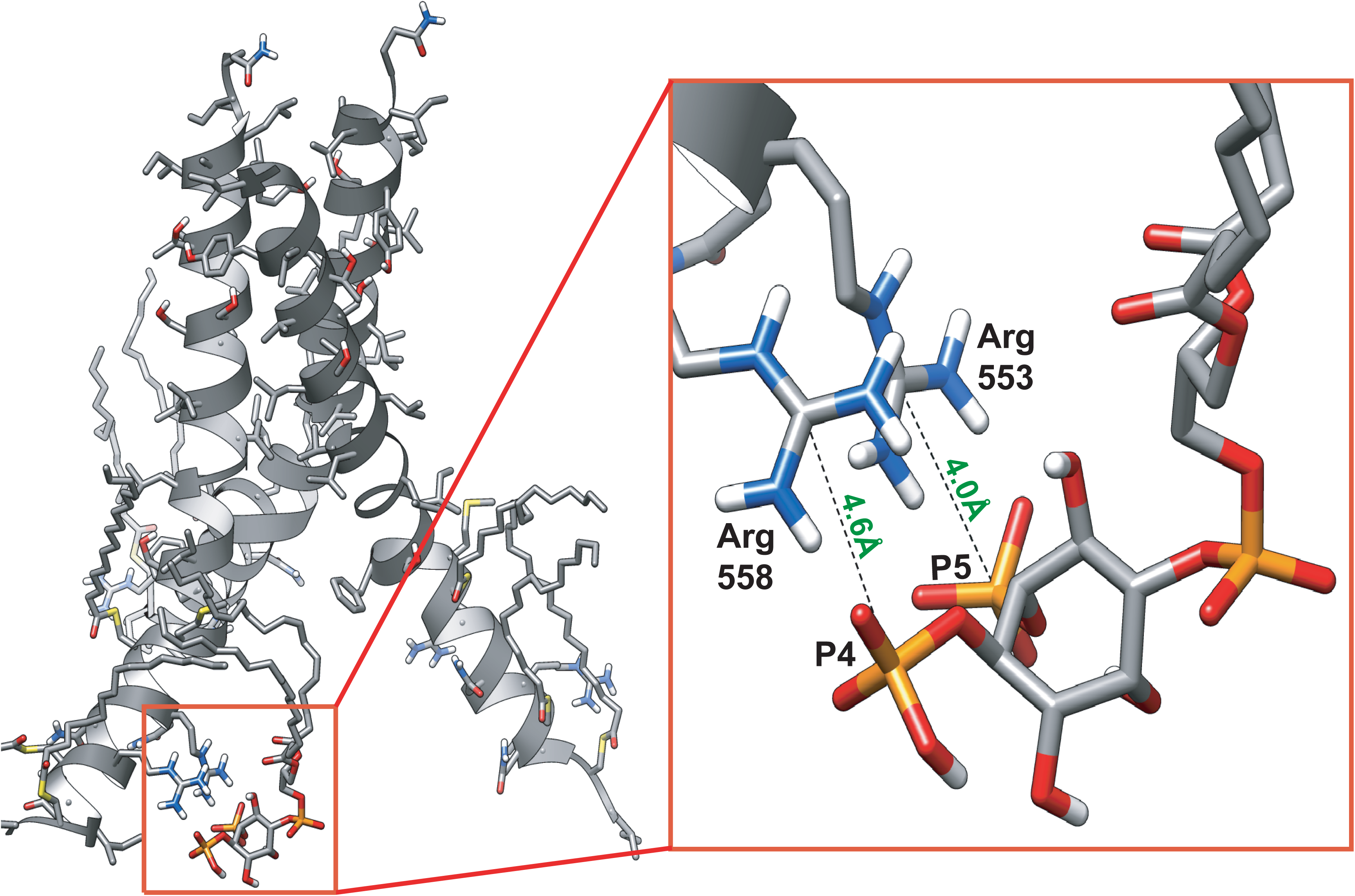
Molecular dynamics simulations reveal direct HA-PIP2 interaction. (A) An example snapshot of HA near a PIP2 molecule pulled from a molecular dynamics simulation showing their close proximity. (B) A magnified view shows the PIP2 headgroups P4 and P5 interacting with the arginines in the cytoplasmic tail domain of HA. Both phosphates are seen to be within 5 Å of Arg 553 and Arg 558. Results are representative of 5 independent replicates.

To characterize and compare interactions between PIP2 and HAwt or HA mutants, RDFs were calculated between the HA and the two phosphate groups on the PIP2, for simulations progressed to 1 μs (Figure 2); RDFs express the degree of enrichment of a moiety as a function of distance from another, where a value of 1 is equal to the average density. Thus, RDF values higher than 1 are enriched, and values lower than 1 are depleted[62]. For the HAwt and the HA mutants that had at least one Arg (553 or 558) present, that Arg was used as the reference (center) for the RDF. For HARREQ (net negative charge in the CTD) and HARREQMAY (net neutral charge in the CTD and no palmitoylation), the Glu-553 and Gln-558 in the CTD were used as references for calculating the RDFs. HAwt has the highest RDF amplitude within 1 nm, as expected (Figure 2A). HAMAY (no palmitoylation and a positively charged CTD) and HARE (net neutral CTD) also have a noticeable peak within 1 nm (Figure 2B-C), while all the other HA mutants are below the average within 1 nm (Figure 2D-H). All the RDFs slowly approach 1 at r>3nm. Those HA mutants that do not have a peak within 1 nm recover to an amplitude of 1 for r>1nm. As a further quantification of interaction, the average RDF from 0 ≤ r ≤ 1 nm was calculated for HAwt and the HA mutants, showing a decreasing trend as Arg-553, Arg-558, Cys-551, Cys-559, and Cys-562 are mutated, with stronger decreases caused by charge-altering mutations, and the strongest decreases observed when both arginines were mutated (Figure 2I).

**Figure 2.**
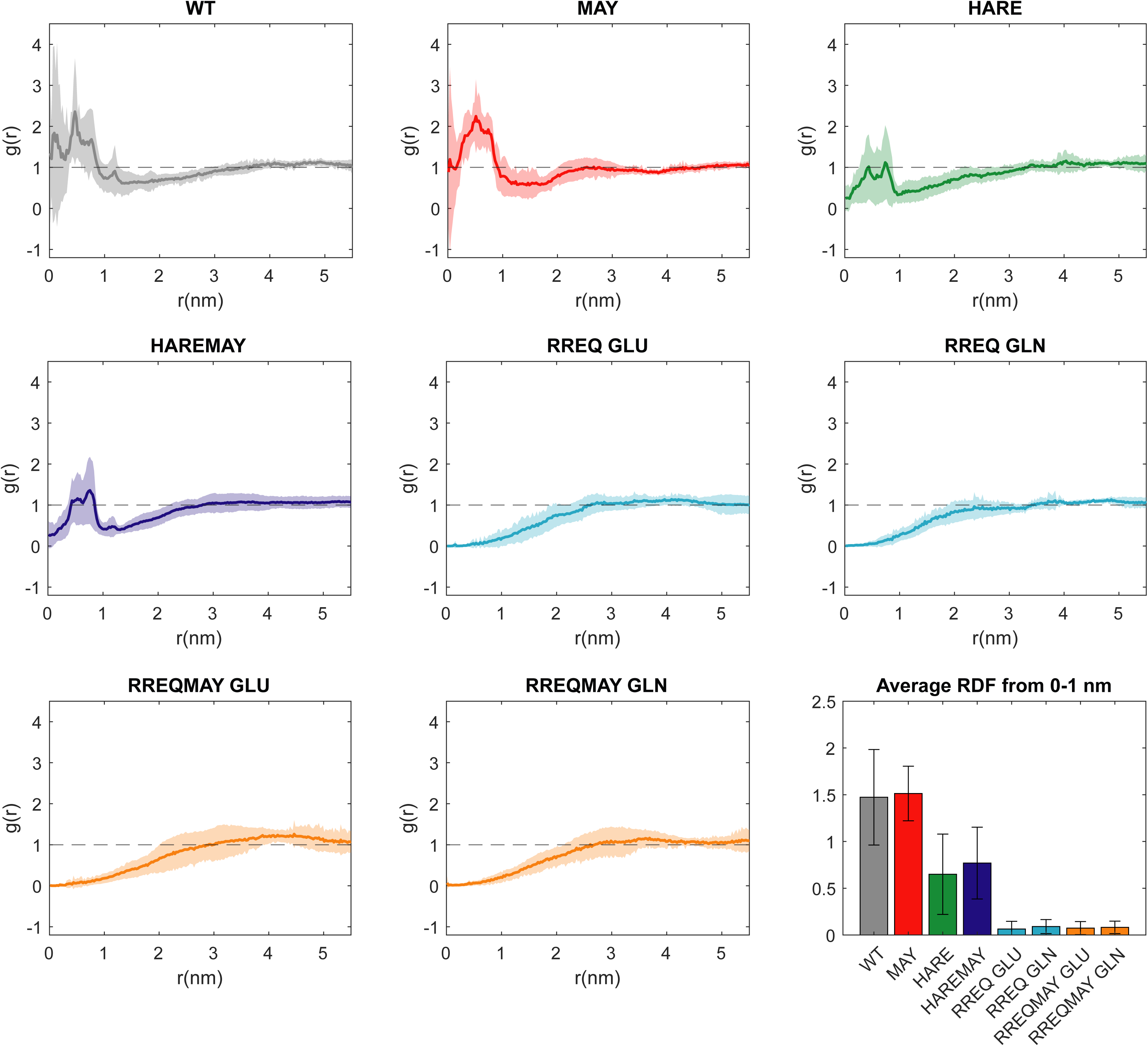
Molecular dynamics simulations show evidence of an electrostatic interaction between PIP2 and the HA CTD. HA and PIP2 in bilayers were simulated as a function of mutations of the HA CTD, and radial distribution functions (RDFs) were calculated between the HA CTD and PIP2. For each HA CTD mutant, 8 simulations were progressed to 2 µs. and the RDF amplitude g(r), is calculated between the CZ atom of arginine and the P4 atom of PIP2 (WT, MAY, HARE, HAREMAY), the CD atom of glutamic acid and the P4 atom of PIP2 (RREQ, RREAMAY), or the CD atom of glutamine and the P4 atom of PIP2 (RREQ, RREQMAY). (A-D) PIP2 headgroups can be found within 1 nm of the cytoplasmic tail domain of HA, HAMAY, HARE, and HAREMAY. (E-H) PIP2 headgroups were not found within 1 nm of the cytoplasmic tail domain of HARREQ or HARREMAY. (I) The average RDF for each HA cytoplasmic tail domain mutant out to 1 nm shows an enrichment of PIP2 for HAwt and HAMAY, a ∼50% depletion of PIP2 for HARE and HAREMAY, and an almost complete repulsion for HARREQ, and HARREQMAY.

Additionally, the clustering of PIP2 with itself was investigated by RDF analysis when an HAwt trimer is nearby in the plasma membrane (Figure S1). It can be seen from the RDF that there is an enrichment, on average, of PIP2 around other PIP2 molecules within 1 nm of the headgroup, showing that PIP2 is clustering with itself in the presence of an HA trimer. PIP2 self-clustering has been observed previously to be mediated by bivalent cations[63, 64], but not in the presence of a viral glycoprotein.

### Palmitoylation of the cytoplasmic tail domain of HA is necessary to maintain alpha helicity

The secondary structure for the CTD was modeled as an alpha helix, which was predicted by AlphaFold[55]. Each of the HA CTD mutants were simulated up to 2μs and MD trajectory positions were calculated for the CTD at the end of the simulation. These trajectories were then analyzed using a custom script (Alex Sodt et al. unpublished) to obtain the probability of the CTD being alpha-helical (Figure 3). This analysis showed that the structure maintains a probability of being alpha helical of ≥75% for residues 555-561. Importantly, removal of palmitoylation (MAY, HAREMAY, and HARREQMAY) reduced the probability for the CTD to remain an alpha helix, compared to HAwt, HARE, and HARREQ (Figure 3). The alpha helix probability is reduced most for residues 562-565, namely the last four amino acids of the CTD, again most strongly for the “MAY” mutants where the three cysteines are mutated to prevent palmitoylation.

**Figure 3.**
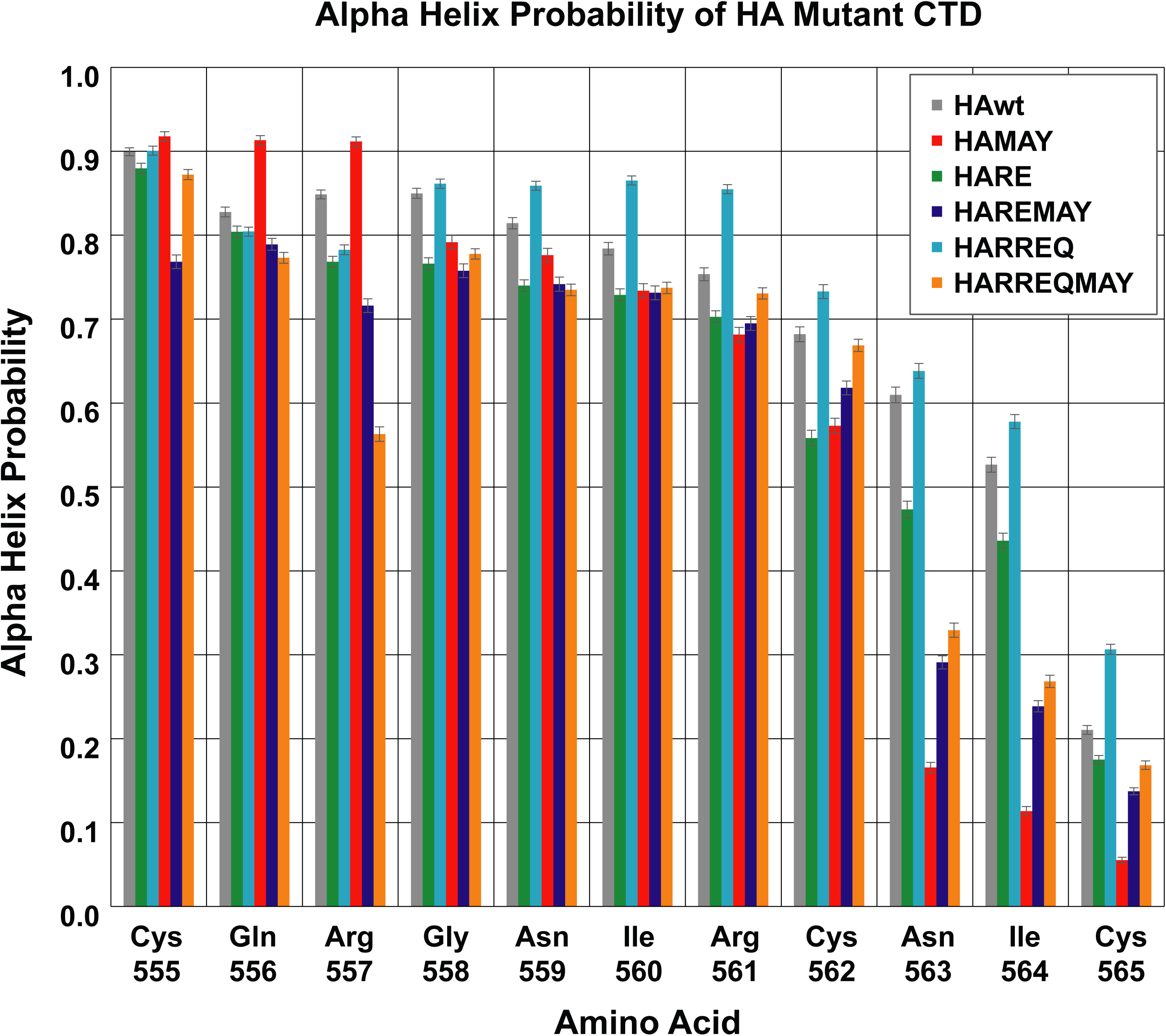
Palmitoylation of the HA CTD affects its secondary structure. The probability of the CTD being an alpha helix was calculated for each HA mutant across all MD sims. A value of 1 signifies that the CTD remained an alpha helix for the duration of the simulations, while 0 signifies that the CTD is no longer an alpha helix. These probabilities were calculated for amino acids 555-565 for each HA CTD mutant, averaged, and graphed. HA CTD mutants that had their palmitoylation sites removed (HAMAY, HAREMAY, HARREQMAY) show an overall reduction in alpha helicity compared to their palmitoylated counterparts (HAwt, HARE, HARREQ) implying that palmitoylation is important for the secondary structure of the HA CTD.

### HA CTD mutations subsequently reduce HA density while increasing PH domain labeling inside PM clusters

Although MD simulations can show us how a single HA trimer can interact with PIP2 on the atomic scale, little can be deduced for HA clustering on the nanoscale. To understand how mutations to the HA- CTD may affect its clustering properties at the plasma membrane, we imaged fluorescently-tagged (Dendra2) wild type HA (HAwt) and its subsequent cytoplasmic tail domain mutants in fixed cells using FPALM-TIRF. We co-expressed PAmKate-tagged PLCδ-PH domain (PH domain) to visualize unbound PIP2 at the plasma membrane.

As previously reported[26], we observed dense clusters of HAwt spanning a diverse range of sizes (∼30-1000 nm diameters) and shapes at the PM (Figure 4A-B). PH domain formed clusters on its own and with HA, while regions along the cell edges were often found enriched with PIP2, in agreement with previous findings (Figure 4A-B)[26, 64–66]. We also observe a distinct lack of HA-PH colocalization within the most dense clusters of HA (Figs. 4B, S2). In comparison, FPALM renders of the HA mutant HAREMAY (with neutral CTD charge and palmitoylation sites removed) showed reduced HA cluster density and more frequent co-clustering with PH domain at the PM (Figure 4C-D).

**Figure 4.**
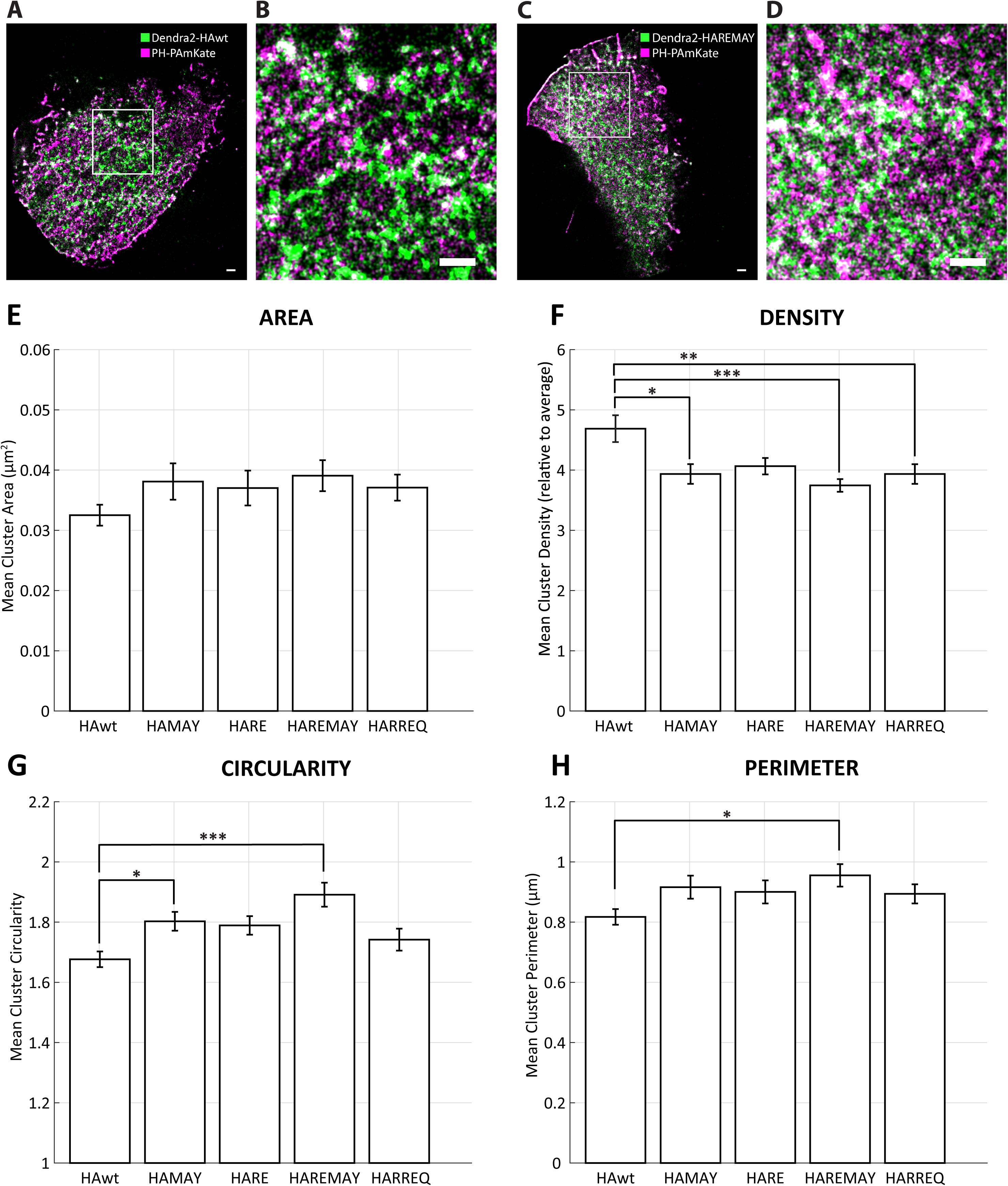
Removal of palmitoylation and net charge neutrality to HA cytoplasmic tail domain drastically changes HA clustering properties. **A-B)** Representative 2-color FPALM-TIRF imaging of HAwt (green) co- labeled with PH domain (magenta) in fixed NIH3T3 cells. Panel B represents a zoomed in region showing areas of colocalization in white. **C-D)** Representative 2-color FPALM-TIRF imaging of HAREMAY (green) co- labeled with PH domain (magenta) in fixed NIH3T3 cells. Panel D represents a zoomed in region showing an increase in colocalization (white) for this cytoplasmic tail domain mutation. All scale bars in panels A-D are 1 μm HA cluster properties were calculated for HAwt and HA cytoplasmic tail domain mutations as follows **E)** area, **F)** density, **G)** circularity, and **H)** perimeter. ANOVA was used for all significance testing where p- value cut offs were indicated as follows, with increasing significance, p<0.05 (*), p<0.01 (**), and p<0.001 (***). Error bars indicate standard error of the mean.

To quantify the spatial distributions of HAwt and HA CTD mutants within clusters, we calculated RDFs for identified clusters of Dendra2-HAwt and Dendra2-HA CTD mutants. In total, we show data for 185 cells with 15,545 total clusters pooled from at least 3 biological replicates. The HAwt RDF was observed to be highest in the centers of the clusters, rising to about 6 times the cell average density, then dropping to about half of the peak density after ∼100 nm (Figure 5-A-D, black data points). RDFs for all HAs are monotonically decreasing, with the most rapid decrease occurring between 70-110 nm, before flattening out and slowly approaching amplitude of 1 (density equal to cell average) as r>200 nm. When compared to HAwt, the central RDF value consistently decreases as mutations are introduced to the CTD. Using a Kruskal-Wallis one-way ANOVA test, we found that RDFs for all of the HA mutants were significantly reduced compared to HAwt for r<100 nm, while maintaining similar RDF shapes overall. HAMAY (palmitoylation sites removed) (Figure 5A), HARE (net neutral charge) (Figure 4B), and HARREQ (net negative charge) (Figure 4D), and were 13.7% ± 6.1% to 18.6% ± 6.7% lower in density than the wild type for r<100 nm. HAREMAY (net neutral charge and removed palmitoylation sites) displays the largest and most significant central reduction 22.1% ± 5.8% (p<0.001) (Figure 4C). Interestingly, we measure a significant increase (10.4% ± 4.7%, p<0.05) in the number of clusters identified per µm^2^ of cell area for HARREQ while the other mutants were not significantly different from HAwt.

**Figure 5.**
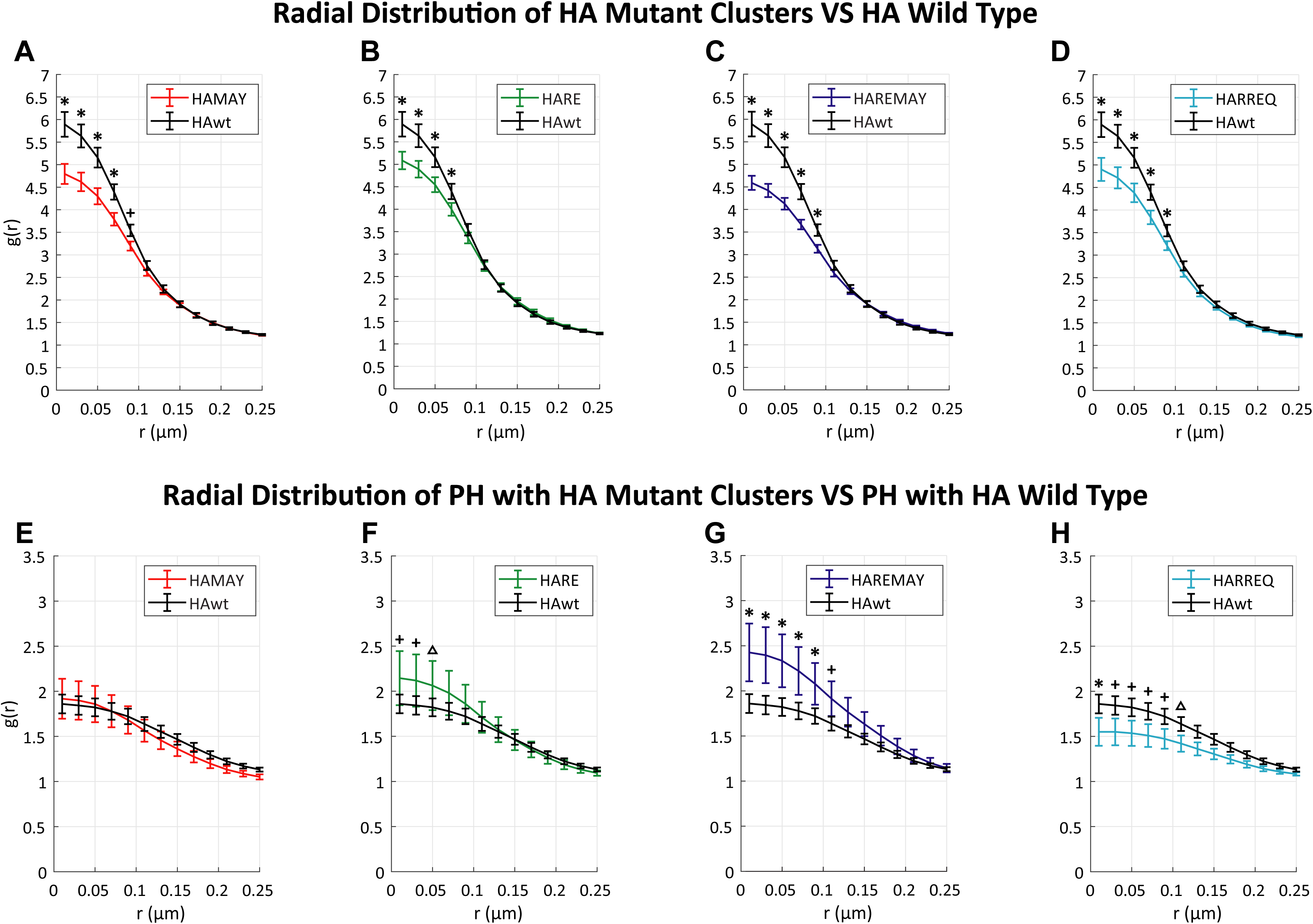
HA cytoplasmic tail domain mutations reduce HA clustering at the plasma membrane but subsequently increase PH domain labeling. **A-D)** Radial distribution functions (RDFs) of HAwt clusters were compared to HA cytoplasmic tail domain mutants HAMAY, HARE, and HAREMAY that show a progressive reduction in cluster density with each mutation. **E-H)** RDFs of PH domain identified within HA clusters show a progressive increase in PH domain with each mutation. A two-way ANOVA test was used to test significance. An asterisk indicates p<0.001, a plus sign indicates p<0.01 and a triangle indicates p<0.05. Error bars indicate standard error of the mean.

To further test the hypothesis of HA-PIP2 interactions through the HA CTD, we calculated RDFs for the PH domain underlying the previously identified HA clusters (Figure 5E-H). The RDFs for PH domain were also maximal at the HA cluster center but dropped to about half of the peak value within 170 nm (compared to 90 nm for HA), suggesting that the PH domain remains enriched past the edges of the HA clusters (Figure 5E-H). PH domain enriched at HAMAY clusters is not significantly changed compared to HAwt (Figure 5E). In contrast, the PH domain RDFS enriched near clusters of HARE, HAREMAY, and HARREQ mutants show significant differences compared to HAwt clusters (Figure 5F-H). PH domain is significantly enriched at the centers of HARE and HAREMAY clusters showing a maximum of 30.5% ± 14.6% and 15.4% ± 15.3% increase respectively from HAwt (Figure 5F-G). Interestingly, PH domain is significantly decreased in the central region for HARREQ clusters at 16.6% ± 11.6% compared to HAwt (Figure 5H).

These differences can be more easily seen if the PAmKate-PH RDF in cells expressing HA-RREQ, where MD simulations showed no interaction between HA and PIP2, is subtracted from all of the other RDFs (Figure 6). The RREQ-subtracted RDFs show enrichment of PH (PIP2) close to HAwt clusters (Figure 6B), and even stronger PH labeling for HAMAY, HARE, and strongest for HAREMAY. Note that this trend in PH-labeling increases as the average HA-PIP2 RDF from MD simulations decrease (Figure 2).

**Figure 6.**
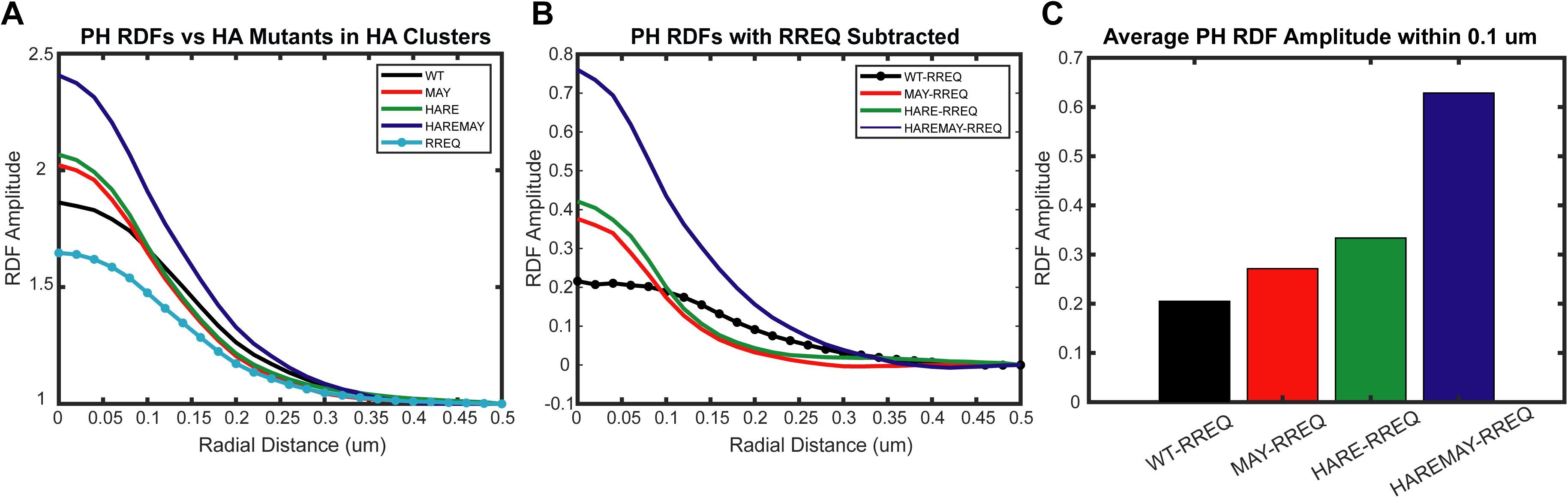
PAmKate-PH Radial Distribution Functions (RDFs) in the vicinity of HA clusters with and without subtraction of the PH RDF near HARREQ clusters. (A) Cluster-based RDFs for PH near HA and HA CTD mutants in fixed cells imaged by FPALM (B) RDFs for PH after subtraction of the RDF for PH near HARREQ clusters (C) Average RDF for PH from r=0 to 100 nm after subtraction of RDF for PH near clusters of HARREQ.

### Removal of palmitoylation and net charge neutrality to the cytoplasmic tail domain of HA drastically changes HA clustering properties

Furthermore, we characterized the HAwt and HA-CTD mutant cluster parameters to elucidate possible changes (Figure 4). There was no significance to the changes measured for the mean cluster area for any of the four CTD mutants compared to HAwt; however, HAREMAY showed the largest increase in area (Figure 4E). Interestingly, HAMAY, HAREMAY, and HARREQ all exhibited significant decreases in the mean cluster density with HAREMAY having the largest decrease (20.1% ± 5.5%, p<.001) (Figure 4F).

HAMAY and HAREMAY had increases in their mean cluster circularities with HAREMAY having the largest increase compared to HAwt (12.8% ± 2.6%, p<.001) (Figure 4G). Cluster perimeters were not significant for all cytoplasmic tail domain mutants with the exception of HAREMAY which was significantly increased (16.9% ± 5.0%, p<0.05) (Figure 4H). The HAREMAY mutant, which has changes to both the charge (net neutral charge) and palmitoylation sites (all removed), was the only measured mutant to exhibit changes in 3 of the 4 properties examined. In each of the 4 properties examined, as well as the RDFs for both HA and PH domain, the HAREMAY mutant exhibits the largest and most significant differences from HAwt, implying a role for electrostatics and palmitoylation in HA clustering mediated by a PIP2 interaction.

### Simultaneous Equilibrium Binding Model for HA, PIP2, and PH-domain

It was observed that cells expressing the highest levels of PH domain unexpectedly showed lower colocalization between PH and HA (Figure S2). Based on published work showing that overexpression of PH domain can lower available PIP2 within the PM[67], we hypothesized that overexpression of PH domain could reduce the PIP2 in the PM which is available to interact with HA. To test this hypothesis, we developed a model of PM levels of HA-PIP2 (HA bound to PIP2) using two simultaneous reversible reactions: the known PIP2-PH binding/unbinding reaction and its equilibrium constant 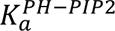,[46] defined as R_2_ in the theoretical model (Supporting Material Eqs. 1-13) and an additional reversible binding between HA and PIP2 (see Supplemental Methods) with an unknown equilibrium constant 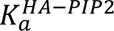, defined as R_1_ in the model (Supporting Material Eqs. 1-13). Note that this model assumes that PIP2 and its complexes equilibrate locally while being spatially correlated with HA as observed experimentally (Figure S3). We then analyzed the spatial overlap fraction (F) of HA with PH domain as a function of the cell-average density of PH-PAmKate (the average number of PH-PAmKate localizations per unit area within a given cell) (Figures 6, S4), and fitted the spatial overlap with the model (Eq. 3 in Supporting Material), which yielded a binding constant R_1_ for HA with PIP2 and a relative binding constant κ for HA-PIP2 binding compared to PH-PIP2: 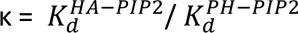. Figure 6 shows the measured F, the fits using the model, and the resulting κ values are shown in Table 2. Although the error bars are large, the κ values are all larger than unity, suggesting that HA-PIP2 binding is stronger than that of a PH domain binding to PIP2.

**Table 2.** Results from nonlinear fit of HA-PIP2-PH competitive binding model (Supporting Material Equations 1-13) predicted HA-PH spatial overlap vs. cell average density of PH. Error bars in the final digit are indicated by the numbers in parentheses following the value. For these results shown, the total [PIP2] was fixed at 1070/μm^2^, R_2_=0.014, and local [HA] (symbol H_0_) was made proportional to total [PH] (symbol K_0_) using H_0_ = 0.475 K_0_. This combination of parameters also minimized the total χ^2^ for the groups combined.

| HA Mutant | Equilib. Const. $R_1$<br>($\mu\text{m}^2/\text{molecule}$) | Relative Equilib. Const.<br>$\kappa$ (relative to PH) | Amplitude $A_0 \times 10^{-4}$<br>(dimensionless) | $\chi^2$ |
| --- | --- | --- | --- | --- |
| HAwt | 0.0747±0.015 | 5.33±1.06 | 5.86±0.18 | 1.14 |
| HAMAY | 0.0512±0.022 | 3.65±1.61 | 6.06±0.63 | 1.63 |
| HARE | 0.0614±0.026 | 4.39±1.85 | 5.85±0.49 | 1.33 |
| HAREMAY | 0.0455±0.019 | 3.25±1.37 | 5.57±0.62 | 1.01 |
| HARREQ | 0.0396±0.016 | 2.83±1.16 | 6.11±0.19 | 0.97 |

Furthermore, the κ values for HA CTD-mutants are smaller than for HAwt, suggesting that changes to the HA-CTD can modulate (and weaken) the binding of HA to PIP2. Notably, the values for HAREMAY and HARREQ are smaller, suggesting substantial inhibition of binding. The binding energy cannot be calculated from the MD simulations because there are insufficient numbers of events where PIP2 unbinds the HA CTD during accessible simulation timescales. Images predicted theoretically using the model reproduce the reduced HA-PH colocalization within the most dense HA clusters, particularly when the interaction between HA and PIP2 is strong (Figure S5), demonstrating further agreement between model and experiment.

## DISCUSSION

Influenza remains an ongoing threat to human health. Better understanding of how Influenza components interact with host cell membranes is needed. Phosphoinositides modulate many cellular membrane functions and have been implicated in the life cycles of other enveloped viruses[31, 67–73]. We first observed the colocalization between PIP2 and HA in fixed cells[26]. We therefore decided to investigate potential interactions between PIP2 and HA with several methods.

### HA-PIP2 Mechanism of Interaction

HA is known to colocalize with PIP2 at the PM[26], however the mechanism for this colocalization is unknown. Previous studies using diffraction-limited and super-resolution methods showed a spatial correlation between PIP2 and HA in fixed cells. We showed previously that in living cells, HA could modulate the diffusion of the PIP2-analog GloPIP[26]. These findings together suggest that there is an interaction between HA and PIP2 at the PM that could be vital to HA clustering. Other studies have shown PIP2-dependent PM trafficking of cellular proteins with polybasic residues and acylation sites[66]. Notably, the CTD of HA contains 1-2 positively charged amino acids along with 3 palmitoylated cystines[20], and these features are highly conserved across influenza strains[25]. We therefore hypothesize that an HA-PIP2 interaction may have two components: 1) an electrostatic interaction between the charged features of the HA CTD and the PIP2 headgroup phosphates; and 2) a hydrophobic interaction between PIP2 and the acyl chains on the cysteines of the HA CTD.

Molecular dynamic (MD) simulations of HAwt with PIP2 in bilayers revealed that the CTD of HA is found close enough to the PIP2 headgroup to have significant electrostatic interaction (Figure 1). To determine the extent of the HA-PIP2 electrostatic interaction, we quantified the radial distribution functions (RDFs) of MD-simulated PIP2 with HA as a function of mutations that altered the charge of the HA CTD (Figure 2). The CTD was mutated in the following ways: 1) removing the palmitoylation sites (HAMAY), 2) reversing one positive amino acid to a negative amino acid resulting in a net neutral charge (HARE), 3) both removing the palmitoylation sites and a net neutral charge (HAREMAY), 4) reversing the CTD charge to net negative (HARREQ), and 5) both removing the palmitoylation sites and a net negative charge (HARREQMAY). By measuring the radial distribution function (RDFs) between the HA CTD mutants and PIP2 headgroups in our MD simulations, we were able to determine that the reversal of the CTD charge to net negative was enough to prevent PIP2 from localizing within 1 nm of the CTD (HARREQ and HARREQMAY) (Figure 2). Removing the palmitoylation sites only (HAMAY) resulted in no appreciable change to the RDFs. There was a small reduction to the average RDF when the CTD net charge was made neutral (HARE and HAREMAY) but there was still evidence of PIP2 within 1 nm of the HA cytoplasmic tail domain in these cases. Only in the case where the charge of the cytoplasmic tail domain was reversed from net positive to net negative did we see a complete repulsion of PIP2 near the cytoplasmic tail domain. This taken together implies a strong electrostatic interaction between the cytoplasmic tail domain of HA and the headgroup of PIP2.

Furthermore, we experimentally quantified RDFs of HAwt and HA CTD mutants with PIP2 using two-color FPALM as a function of the local levels of PH domain (Figure 5). We imaged HAwt and the HA CTD mutations HAMAY, HARE, HAREMAY, and HARREQ in fixed cells each co-expressed with PH domain to visualize PIP2. Our fluorescently tagged HARREQMAY mutation did not express well in cells and we were unable to receive reliable data for this study. Using MD and FPALM results together and comparing HAwt to HARREQ, we see a significant decrease in the RDF by FPALM and a reduction of RDF below 1 in the MD simulations. This suggests that there is strong disruption of the interaction and exclusion of the PIP2 from the HA clusters, likely due to the electrostatic repulsion between the negatively charged HARREQ CTD and the negative headgroup of PIP2. In the FPALM experiments, the RDF for PH domain found within HA clusters is not as depleted as in the MD simulations. By mutating the CTD to be net negative (HARREQ) we would expect to find no PIP2 in HA clusters as predicted by electrostatics alone. However, we observed a decrease but still a non-zero density of HA near PH clusters observed by FPALM (Figure 5). This suggests that there are other factors keeping PIP2 near HA, which we speculate could include 1) HA interactions with other proteins which also interact with PIP2[26], 2) confinement due to the actin cytoskeleton[66, 74], 3) presence of ions[63] or 4) other unidentified interactions.

To better isolate the changes in HA-PIP2 colocalization due to changes in the HA-CTD, we used the finding from the MD simulations that HARREQ repels PIP2 and therefore has no attraction to it. Thus, any experimental colocalization observed between HARREQ and PIP2 must be due to other factors. Therefore, the experimental RDF for HARREQ was subtracted from the RDFs for HAwt and HA mutants (Figure 6).

These subtracted RDFs show a trend of increasing HA-PH colocalization as the CTD mutations increasingly disrupt the HA-PIP2 interaction (Figure 6). Considering that PH can only bind free PIP2, if HA is strongly interacting with PIP2, it will locally deplete free PIP2 from the PM, especially in or near dense HA clusters. In support of this explanation, we observed less PH labeling near high-density HA clusters (Figure S2) and less PH labeling in HAwt clusters than in HA mutant clusters (Figure 5). MD simulations show at the molecular scale how mutating the CTD of HA modifies the electrostatic interaction between HA and PIP2, corroborating a molecular driving force for the larger scale (10s or nanometer) two-color FPALM experiments.

MD simulations have also illuminated how removal of the palmitoylation sites affects the interaction between HA and PIP2 and on the HA-CTD structure itself. RDFs from MD simulations did not show much change in overall PIP2 enrichment near HA when palmitoylation was removed (Figure 2). We therefore hypothesized that removal of palmitoylation sites might alter the secondary structure of the HA CTD rather than its direct attraction to PIP2. Thus, the alpha helicity was calculated for the CTD of each HA mutant (Figure 3). When comparing HAwt vs. HAMAY, HARE vs. HAREMAY, and HARREQ vs. HARREQMAY, all the non-acylated (MAY) mutants have a lower alpha helix population than their acylated counterparts, especially for residues beyond Cys562. Comparing HARREQ to HAwt, almost every position in the CTD of HARREQ has a higher probability of staying an alpha helix than HAwt. This could be because there is no electrostatic interaction happening between the CTD of RREQ and PIP2, which suggests that PIP2 is influencing the secondary structure of HA. Based on the MD simulations, there is typically at least one PIP2 within a few nanometers of each HAwt monomer, and this nearby PIP2 can cause the CTD to be pulled in different directions, leading to unraveling of the alpha helicity over time, in the absence of the acylated cysteines.

Two-color FPALM imaging of PAmKate-PH coexpressed with HAMAY or HAREMAY and compared to HAwt complements the molecular simulations. The RDF for the PH in HA clusters shows modest differences between HAwt and HAMAY (Figure 5) suggesting that the palmitoylated sites do not have a strong effect in the HA-PIP2 interaction. When comparing the RDF of HAREMAY and HAwt (Figure 5), a significant increase in the amount of free PIP2 being labeled by PH domain is observed (Kruksal-Wallis one-way ANOVA test for each mutant compared to HAwt). Furthermore, the RDFs for HAREMAY and HAwt remain significantly different out to 110nm (Figure 5G), compared to 50nm comparing HARE with HAwt (Figure 5F), suggesting larger changes in HA-PH colocalization for HAREMAY than for HARE. Differences in HA-PIP2 interactions caused simply by removal of acyl groups (i.e. MAY) may not be strong enough alone to yield significance in the cellular images, while removal of one charge did yield some significance at short range, and removal of both charge and acyl group yielded significance over a larger range of distances. Because HARREQMAY could not be successfully imaged, we cannot compare FPALM quantification for HARREQ with HARREQMAY.

We have been able to quantify the HA-PIP2 interaction by using both MD simulations in parallel with experimental single molecule (FPALM) imaging. The information provided by these two methods is complementary: in MD simulations we directly observe PIP2 in a model PM, and directly observe the molecular interactions between HA and PIP2. On the experimental side, we do not directly observe PIP2, but rather use the PIP2-binding protein PH-PLC*δ*1 to observe free PIP2 in the PM. Because the biosensor we are using only binds to free PIP2, we will not see PIP2 bound to HA or other molecules. Thus, when HA is interacting with PIP2, PH-PLC*δ*1 cannot interact with that same PIP2. Conversely, PIP2 bound to PH-PLCδ1 presumably cannot interact with HA. It is possible that HA could interact with PH-PLC*δ*1[74], but such interactions would predict higher HA-PH colocalization at higher PH levels, which is opposite to our experimental observations (Figure 7).

**Figure 7.**
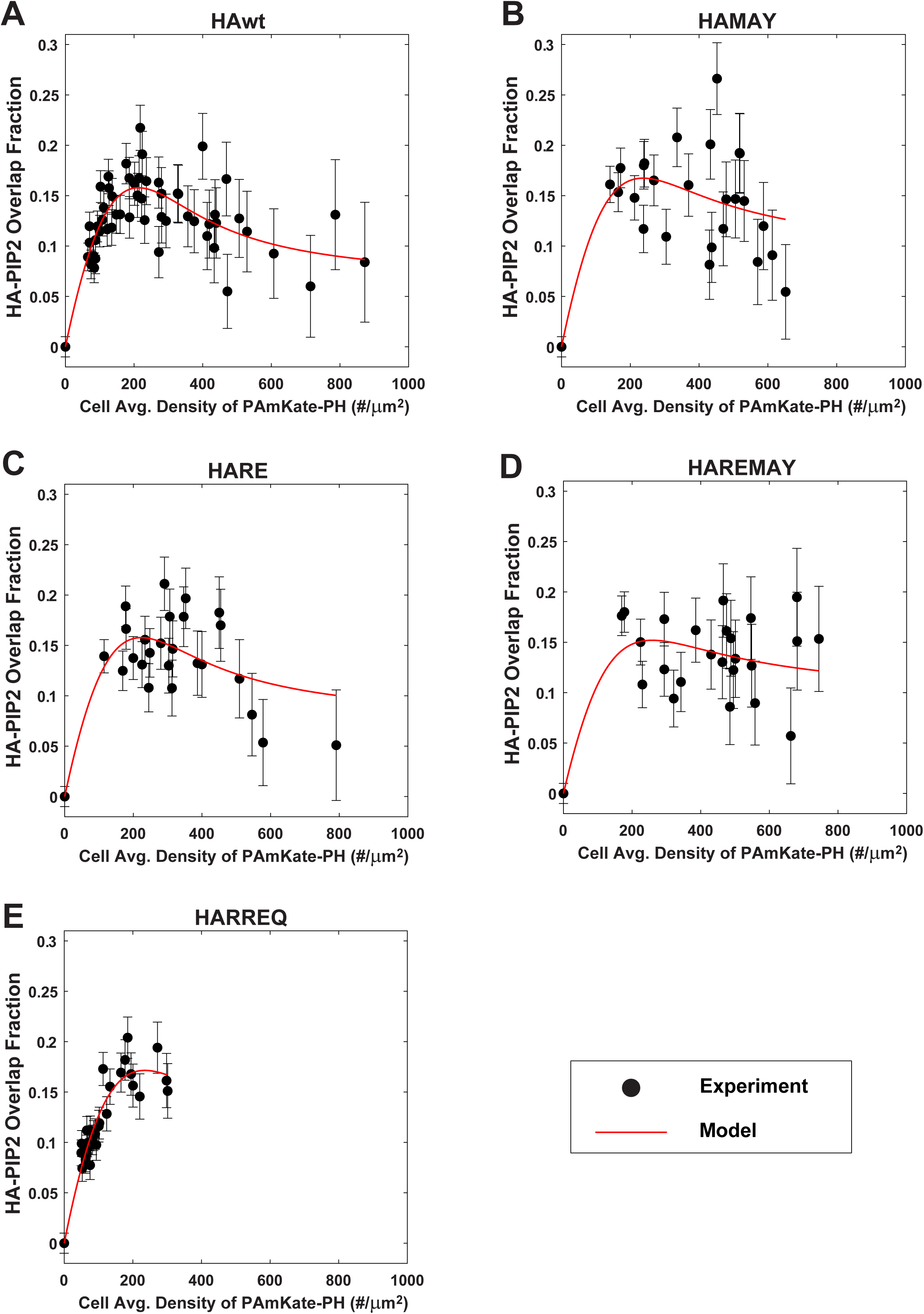
**C**ompetitive binding model for HA, PIP2, and PH fits experimental colocalization of HA and PH imaged by FPALM. The spatial overlap fraction of HA and PIP2 is plotted as a function of cell average density of PH-PAmKate for **A)** HAwt, **B)** HARE, **C)** HAMAY, **D)** HAREMAY, and **E)** HARREQ. Experimental data (black dots with error bars equal to standard error of the mean) are shown together with the competitive binding model fit (red line).

Interpretation of these findings has been greatly helped by a model of competitive binding between HA, PIP2, and PH-PLC*δ*1 (See Supporting Material Eqs. 1-13). It was observed that the cells expressing the highest levels of PH-PAmKate showed lower colocalization between HA-Dendra2 and PH-PAmKate (Figure 5, 7, S2). Based on previous work showing that overexpression of PH domain can lower available PIP2 within the PM[28, 75, 76], we hypothesized that overexpression of PH domain could reduce the free PIP2 in the PM which is available to interact with HA. To test this hypothesis, we developed a model of two simultaneous reversible reactions: the known PIP2-PH binding/unbinding reaction with its equilibrium constant K_aPH-PIP2_[46], and an additional reversible binding between HA and PIP2 with an unknown equilibrium constant K_aHA-PIP2_ (see Supporting Material). As noted above, the model assumes that PIP2 and its complexes equilibrate locally while being spatially segregated by other unknown factors. We then analyzed the spatial overlap fraction (F) of HA with PH in FPALM datasets as a function of the cell-average density of PH-PAmKate (the average number of PH-PAmKate localizations per unit area within a given cell), and fitted the spatial overlap with the model (Supporting Material Eq. 3), which yielded κ = K_aHA-PIP2_/ K_aPH- PIP2_. Figure 7 shows the measured F, the fits using the model, and the resulting κ values. Although the error bars are large, the κ values are all larger than unity, suggesting that HA-PIP2 binding to PIP2 is weaker than for a PH domain, although comparable in magnitude for HAwt. Furthermore, the κ values for HA CTD- mutants are smaller than for HAwt (Table 2), suggesting that changes to the HA-CTD can modulate the binding of HA to PIP2. Since these mutations do not completely abolish HA-PIP2 colocalization (Figure 5), there must be other factors controlling their colocalization, independent of their direct interaction.

Importantly, the chemical model explains the reduction of HA-PH colocalization at high levels of HA (Figure S2) or PH (Figure 7), confirms disruption of HA-PIP2 interactions when the HA CTD is mutated (Figure 7), and suggests that HA can compete with other cellular proteins for binding of PIP2, which could modulate cellular functions if HA is present at high enough concentrations. The model was used to generate simulated imaging results (Figure S5) which qualitatively reproduce the experimentally observed patterns in HA-PH colocalization (Figures 5, S2). A cartoon summary of the essential aspects of the interaction model is shown in Figure 8.

**Figure 8.**
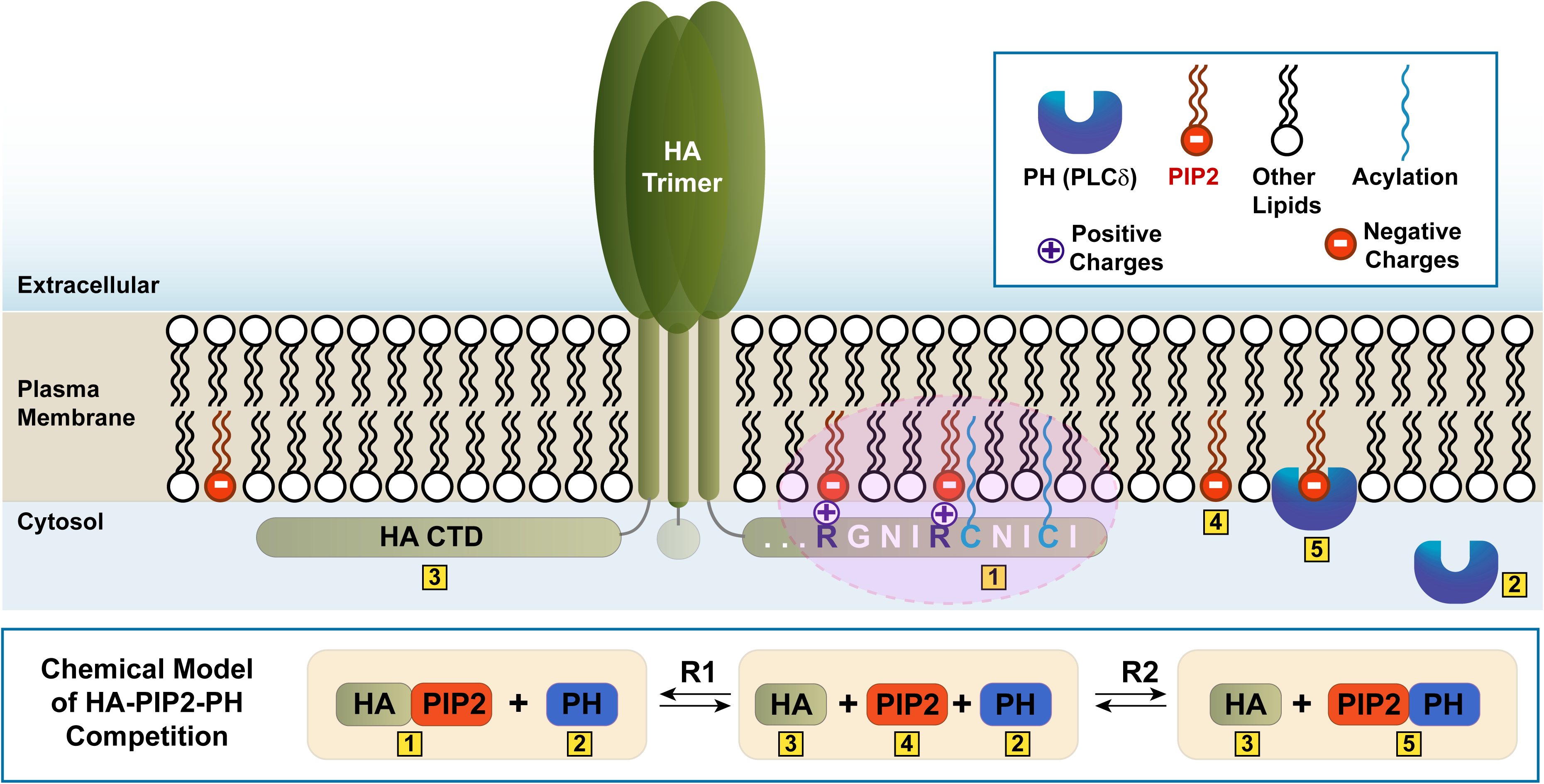
Model of proposed PIP2 interaction with the HA cytoplasmic tail domain. The model correlates the experimental observations of chemical equilibrium between free PIP2 and PIP2 bound to HA or PIP2 bound to PH domain. Cytosol - Light aqua blue region; Plasma membrane - Beige region; Extracellular - Cream region; HA - Ectodomain : Green trimer, Transmembrane domain - Green vertical tube, Cytoplasmic tail domain: Green horizontal tube with residues ; Arginine - Purple R with a (+) charge; Cystine - Cyan C; Acylation of C – Wavy line in cyan; PIP2 - Black squiggly lines with (-) charged red headgroup; Lipids - Black squiggly lines with white headgroup; PH - Blue purple U ; Box-1 : PIP2 interaction with HA cytoplasmic tail domain (Purple shaded region), Box-2 : free PH, Box-3 : HA ; Box-4 : free PIP2 ; Box-5 : PIP2 interaction with PH; R1 - Equilibrium of Box-1 and Box-2 with Box-2, Box-3, and Box-4; R2 - Equilibrium of Box-2, Box-3, and Box-4 with Box-3 and Box-5.

### Role of HA-PIP2 Interactions in PIP2(PH) Clustering

PIP2 is a minor lipid component located in the cytoplasmic leaflet of the host cell PM, where it makes up about 1% of the total lipids, yet performs numerous cellular functions related to membrane organization and signaling[67, 69, 70, 77]. Therefore, the processes that control the PIP2 distribution in cell membranes are important for many reasons[66, 68]. A currently unsettled question is whether PIP2 forms clusters in live cell membranes. Hammond et al proposed several membrane models for PIP2-protein interactions, including: the platform, selfish synthesis, and megapool models, which describe ways membrane proteins, lipid kinases, and phosphoinositides can modulate biological membrane organization[78]. Our recent studies show a heterogeneous distribution of fluorescently-labeled PIP2 and PH-labeled PIP2, as well as PIP2 coclustering with viral spike proteins from influenza A[26], influenza B[25], and SARS-CoV2[79]. In addition to the importance of understanding PIP2 distribution in living cells, the involvement of PIPs in viral life cycles provides another compelling reason for determining whether PIP2 does cluster in live cells, and for understanding the mechanism by which such clusters arise. We have a unique position to address this question because we can combine atomistic MD studies of PIP2 with experimental single molecule studies of cellular proteins and free PIP2 observed by PIP2 binding proteins.

In the MD simulations, PIP2 is seen to form small clusters with itself in the PM consisting of 3-4 PIP2s that are scattered throughout the PM (Figure S1). Experimental results showed previously that PIP2 self-clusters in the presence of cations[63]. The cations, especially bivalent cations, can bridge the PIP2 headgroups together which can lead to longer-range structures[80]. In our simulations, the levels of PIP2 were 2-5x higher than what is typically found in a mammalian PM, which increases the rate of PIP2 interactions with HA, but is also consistent with PIP2 levels observed experimentally in clusters[25, 26, 64, 65, 79]. The PIP2 clusters observed in MD simulations are often formed several nanometers from HA (Figure S1), suggesting that they are likely to form in the absence of HA, as has been observed experimentally[26, 63, 64].

We hypothesize that when the electrostatic mechanism of the HA-PIP2 interaction is weakened, this causes a decrease in the amount of PIP2 binding to HA, leading to additional free PIP2 in and near HA clusters, which can then be bound by PH domain. Of course, for HA and PH domain to colocalize, the enrichment of PIP2 in and near HA clusters must be maintained, presumably by attractive interactions between HA and PIP2, restriction of diffusion, uncharacterized interactions with other proteins or lipids, or other mechanisms. Looking at the FPALM images of HA and PH domain (Figure 4), the density of PH-labeled PIP2 increases for the mutants HAMAY and HAREMAY, compared to HAwt. This suggests there is more free PIP2 around the identified (mutant HA) clusters. Since these mutations weaken the electrostatic interaction, fewer PIP2s will be interacting with an average HA, but as long as there is sufficient attraction between the HA and PIP2, the PIP2 can be enriched around HA clusters. These findings are also explained quantitatively by the model of HA-PIP2-PH competition (Figure 7). Together, these results support the hypothesis that PIP2 can interact directly with the HA CTD. Since many other viral proteins contain similar sequence features[25], we anticipate that these interactions could be important for understanding mechanisms of infection of many types of viruses.

Although utilizing MD simulations along with SMLM techniques is powerful, there are still some limitations to these techniques that need to be addressed to better answer the questions regarding protein-lipid interactions. MD simulations are limited by the size of the system and length of simulation. In our MD simulations we have 1 HA trimer embedded in a small portion of a simulated mammalian PM and reach a maximum timescale of a few μs. In an ideal situation, we would simulate the PM of a cell with at least two HA proteins to investigate HA clustering on timescales long enough to allow HA-HA distances to change significantly and PIP2 to bind and unbind repeatedly. Although in principle this type of MD simulation is achievable, it is beyond the scope of this work. Concerning our FPALM experiments, probes that can visualize PIP2 directly (without perturbing its distribution or binding to other molecules) are strongly needed[74, 81, 82]. In consideration of this need, we realized that using our competitive binding model with a known equilibrium constant R_1_ for one PIP2-binding protein (HA) and R_2_ for PH-PIP2 binding, respectively, local observations of [HA] and [PH] can be used to calculate local [PIP2]. This approach could provide much needed quantification relating to models of phosphoinositide distribution and normal cell function. While the presence of overexpressed HA and PH domain will likely alter the PIP2 distribution within the PM, in principle similar methods could be devised using combinations of proteins with different PIP2 binding constants and quite low expression levels to minimize perturbation of cell function and obtain the same kind of information.

### Role of HA-PIP2 Interactions in HA Clustering

It was observed previously that HA cluster properties varied as a function of the level of PH-domain- labeled PIP2[26]. We showed in the previous section that mutations of the arginines and cysteines in the HA CTD altered HA colocalization with PH-labeled PIP2 (Figures 5,6) and that these same mutations disrupted direction interaction between HA and PIP2 (Figures 5,6). Thus, we wanted to see if these interactions affected HA clustering. We observed from experimental FPALM data (Figure 5) that the same mutations that affect HA-PIP2 interactions also influence how HA is clustering with itself. Looking at the cluster properties (Figure 4) there is a significant difference in the density of the identified HA clusters comparing HAwt with HAMAY, HARREQ, and HAREMAY. Between HAwt and all the CTD mutants, there is a decrease in the density of the clusters (Figure 4F). One possible explanation for the differences between HAMAY and HAwt and between HARE and HAREMAY, is that the removal of the palmitoylation decreases HA CTD anchoring in the membrane. The disruption of this anchoring could decrease the HA-PIP2 interaction by allowing the CTD to be moved further away from the cytoplasmic leaflet, which would weaken the electrostatic interaction between HA and PIP2. By decreasing the interaction between HA and PIP2, this leads to decreases in the HAMAY and HAREMAY cluster density (Figure 4F). Considering electrostatic changes to the HA CTD, HARREQ keeps the palmitoylation but reverses the charge of the CTD (Table 1). This is expected to cause the HA CTD to repel PIP2 and is confirmed by the MD simulations (Figure 2). PIP2 is also known to mediate trafficking of membrane proteins to the PM[66], predicting reduced HA density in the PM for HA mutants which interacted less strongly with PIP2, which is confirmed by FPALM imaging (Figures 4F,5D,6A). Thus, these findings show that mutations which disrupt HA-PIP2 interaction also reduce HA cluster density.

Despite the importance of HA clustering to viral infection and human health[4, 11, 17, 83–85] the mechanism of HA clustering and the factors determining HA cluster stability remain unclear[29, 86, 87]. Our results show a significant dependence of HA clustering on HA-CTD tail charge and palmitoylation.

Modifications to the tail clearly caused disruption of clustering, with varying effects on the amount of local free PIP2 and thus PH domain labeling of the PM. Since certain changes to the HA-CTD reduced the relative densities of clusters, those CTD residues must play some role in the clustering of HA at the membrane, indicating it is unlikely that HA is clustered exclusively due to targeting certain membrane regions through the transmembrane domain[88].

### Broadening the Mechanisms of HA-PIP2 Interaction to other Viral Protein-Lipid Interactions

Having demonstrated and quantified the HA-PIP2 protein-lipid interaction, we now ask whether this relationship occurs between other viral spike proteins and PIP2. HA is known to have a highly conserved CTD[89, 90] which we show plays a key role in the HA-PIP2 interaction[25]. It has been shown that other viral spike proteins have similar motifs that are conserved like we see in HA[25]. By identifying the amino acid motifs that remain constant in these highly mutating proteins, these potential sites of interaction with PIP2 or similar lipids could serve as antiviral targets.

We hypothesize that the CTDs of other viral spike proteins can interact directly with PIP2 using the same mechanisms identified above. The CTDs of HA from both Influenza A and Influenza B, including avian influenza A strains, contain highly conserved sequence patterns with positively charged amino acids and cysteines[25], whose propensity for acylation has been studied previously[12, 90]. We showed with MD simulations that Influenza B interacts directly with PIP2[25]. FPALM showed that PIP2 colocalizes with not only IAV HA but also Influenza B HA[25] and the spike protein of SARS-CoV-2[79]. Further work on other viral spike proteins is needed to experimentally test how far this pattern extends.

Considering viral proteins in general, Chukkapalli et al. showed interaction of PIP2 and human immunodeficiency virus type 1 (HIV-1) Gag protein[40]. Although Gag is not considered a spike protein, it is a viral structural protein required for HIV infection. Gag is known to have a patch of conserved residues that contribute to the binding of the protein to the membrane that are myristoylated and palmitoylated. By mutating the MA sequence Chukkapalli et al was able to show that Gag requires certain amino acids in the MA domain to interact with PIP2[40, 41]. Additionally, Ebola virus matrix protein (VP40) has been shown to interact with PIP2[38]. VP40 is not a viral spike protein; it forms a shell underneath the lipid bilayer of the virus, and does not contain a CTD. Johnson et al showed groups of lysines that would interact with PIP2, suggesting an electrostatic interaction[38]. We have shown previously using FPALM that IAV matrix protein M1 coclusters with PIP2[79]. M1 is known to form a layer on the inner leaflet of IAV, similar to VP40 in Ebola virus, and has regions of lysines and arginines. The colocalization of M1 and PIP2 at the plasma membrane of cells suggests there may be an electrostatic interaction present between the two.

PIP2 has been shown to interact with many viral proteins as mentioned above, but it is not the only phosphoinositide present in cells. Phosphoinositides are known for regulating cell signaling and membrane dynamics and could play a role viral protein organization through the same mechanisms we identified for HA-PIP2. We would like to investigate other phosphoinositide species through MD simulations and single molecule (FPALM) imaging as there is a need to research more than PIP2 in the PM[91]. Additionally, there are anionic lipids other than PIPs in our host cells that could also be interacting with viral proteins the same way as HA-PIP2. Using our findings about how HA and PIP2 interact, we can explore other viral protein and lipid interactions to better understand the life cycle of viruses.

We have shown evidence that conserved viral protein features (basic amino acids in proximity to acylated cysteines) have the potential to interact with phosphoinositides. In fact, much of this work was inspired by the fact that normal cellular proteins use similar features for phosphoinositide interactions in order to traffic to the plasma membrane[28]. We also propose that these protein-lipid interactions could provide new anti-viral drug targets. MD simulations can be utilized to identify lipid binding sites on membrane proteins for drug design[91]. What remains to be determined is how to disrupt the interactions of viral proteins with phosphoinositides while leaving normal cell functions undisrupted. Considering phosphoinositides, greater understanding of the specificity of these interactions, and of the cellular components which regulate the relevant phosphoinositide pools, is needed so that viral functions can be specifically attacked while leaving cellular functions intact.

## Supporting information

Winski et al Supporting Material

## Author Contributions

Conceptualization – STH, AS, JZ

Methodology – DW, STH, AS

Resources – DW, STH, AS, JZ, HW

Writing (Review and Editing) – DW, STH, CW, SS, JW, AS, MP, HW, JZ

Project Administration – STH

Funding Acquisition – DW, STH, AS

Supervision – STH

Investigation – DW, STH, PR, MP, JW, SS, CW

Software – DW, STH

Formal Analysis – DW, STH, CW, SS, PR

Visualization – DW, STH, CW, SS

Writing (Original Draft) – MP, JW, DW, STH, CW, SS, HW

## Acknowledgments

The authors would like to thank UMaine ARCSIM for helping with and use of the UMaine supercomputer, and Starr Campbell, Pat Byard, Tim Campbell, and Mariana Haletska for technical assistance as well as Dean Astumian, Brandon M. Aho, and Komala Shivanna for valuable discussions. This work was funded by the National Institutes of Health awards R15GM155811 (PI: Hess), R15GM139070 (PI: Hess), and R15GM116002 (PI: Hess), by the Maine Technological Asset Fund (MTAF 1106 and 2061, PI: Hess), the University of Maine Office of the Vice President for Research, and the Maine Economic Improvement Fund. This work used Expanse GPU at SDSU through allocation number BIO230058 (PI: Hess) and BIO200072 (PI: Hess) from the Advanced Cyberinfrastructure Coordination Ecosystem: Services & Support (ACCESS) program, which is supported by National Science Foundation grants #2138259, #2138286, #2138307, #2137603, and #2138296.

