## Supplementary material for "The Influenza Hemagglutinin Cytoplasmic Tail Domain Interacts with Phosphatidylinositol 4,5-bisphosphate": Winski et al Supporting Material: Winski et al Supporting Material 2026Aug24.pdf

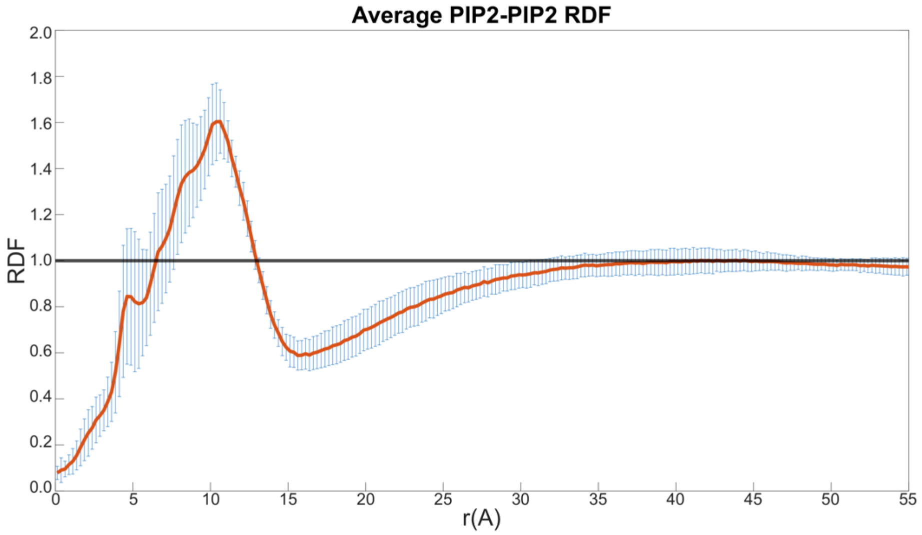


**Figure S1. Quantification of PIP2 Radial Distribution Function.** The radial distribution function (RDF) was calculated from molecular dynamics trajectories of PIP2 molecules simulated in a bilayer with an HAwt trimer. The distance r was measured between the P4 atom of each PIP2. There is an enrichment of PIP2 for r≤1.5 nm, indicating that PIP2 is clustering with itself in the presence of HA.

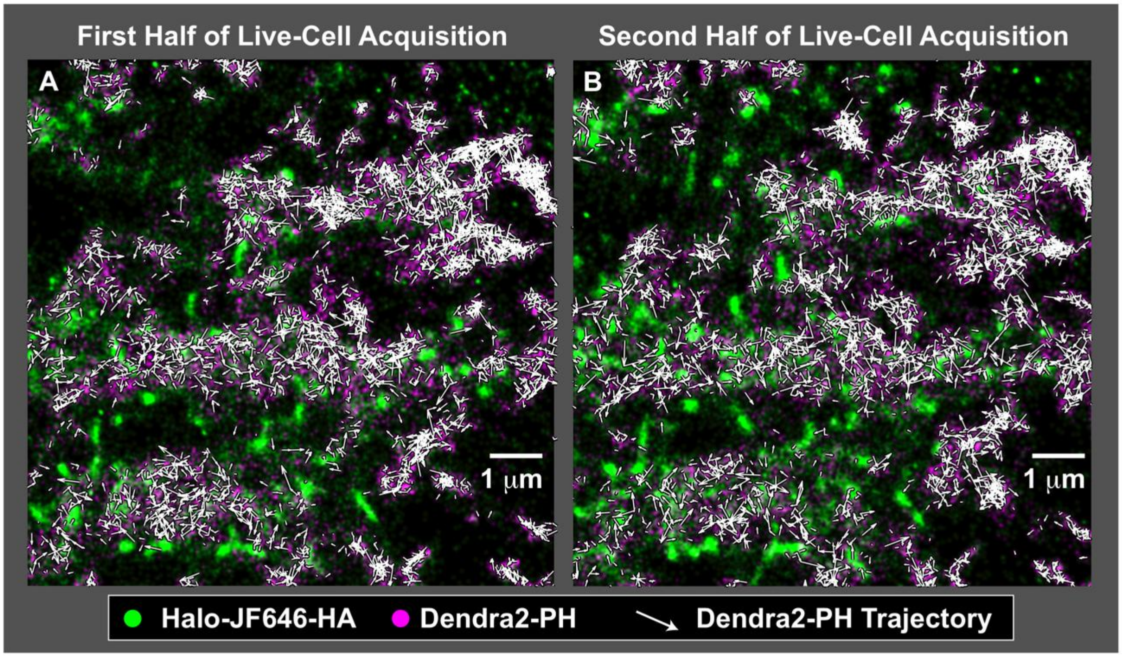


**Figure S2. Live-cell FPALM imaging of Halo-JF646-labeled HA and Dendra2-PH in NIH3T3 cells shows dynamics of PH domain in the vicinity of HA clusters.** (A) First 100 seconds of acquisition (B) Next 100 seconds of acquisition. Note the exclusion of PH from HA clusters with high density. At more modest HA cluster density, colocalization between HA and PH is observed. Note also that HA and PH have a dynamic distribution, as can be seen by comparing panels A and B, which show the same ROI within the cell.

**Theoretical Model of Simultaneous Equilibrium of HA, PIP2, and PH-domain** We consider the putative reversible binding between HA and free PIP2 as an equilibrium reaction with equilibrium constant $R_1$

$$A + P \overset{R_1}{\rightleftharpoons} B \tag{1}$$

and the reversible binding between the PAmKate-PH-domain and free PIP2 with equilibrium constant $R_2$:

$$K + P \overset{R_2}{\rightleftharpoons} Q. \tag{2}$$

where the following symbols for the local concentrations (denoted by square brackets) of the five molecular species are: A = [Free HA], P = [Free PIP2], B = [HA-PIP2], K = [PAmKate-PH], Q = [PAmKate-PH-PIP2].

From the super-resolution microscopy datasets, we can estimate the average of Q over the cell, and measure F, the fraction of pixels which are high in both PAmKate and HA. This can be calculated as the fraction of membrane-associated PAmKate-PH ($Q/K_0$) times $H_0$, the total local concentration of HA:

$$F = \frac{H_0 Q}{K_0}. \tag{3}$$

Where $K_0$ is total PH concentration. Then, from the equilibrium reaction (Eq. 1)

$$R_1 = \frac{B}{A \cdot P} \tag{4}$$

$$R_2 = \frac{Q}{K \cdot P} \tag{5}$$

$$B = APR_1 \tag{6}$$

By conservation of HA molecules, the sum of free HA (A) and PIP2-bound HA (B) equals $H_0$:

$$A + B = A + APR_1 = A(1 + PR_1) = H_0 \tag{7}$$

$$P + B + Q = P_0 \tag{8}$$

$$K + Q = K_0 \tag{9}$$

Combining ,

$$aP^3 + bP^2 + cP + d = 0 \tag{10a}$$

With a=$R_1R_2$, b=$R_1R_2H_0 + R_1R_2K_0 - R_1R_2P_0 + R_1 + R_2$, (10b)
c=$R_1H_0 + R_2K_0 - R_1P_0 - R_2P_0 + 1$, and d=$-P_0$.

This has one real analytical solution (and two complex solutions not shown):

$$P = D_1 + D_2 + D_3 \tag{11}$$

with

$$D_1 = \frac{\left[\sqrt{(-27a^2d+9abc-2b^3)^2+4(3ac-b^2)^3}-27a^2d+9abc-2b^3\right]^{1/3}}{3a\sqrt[3]{2}} \tag{12a}$$

$$D_2 = \frac{-\sqrt[3]{2}\,[3ac-b^2]}{3a\left[\sqrt{(-27a^2d+9abc-2b^3)^2+4(3ac-b^2)^3}-27a^2d+9abc-2b^3\right]^{1/3}} \tag{12b}$$

$$D_3 = -\frac{b}{3a} \tag{12c}$$

Which provides values of P, which can then be used to calculate K, Q, A, and B, and finally F. We use the proportionality between HA and PH observed experimentally (Figure S3) to further constrain the model to relate $H_0$ and $K_0$:

$$H_0 = c_0 + c_1 K_0 \tag{13}$$

*Experimentally-Derived Parameters Used for Model.* To constrain the parameters within the model, values determined by or derived from experiment were used as much as possible. The estimated total PIP2 concentration ($P_0$) was determined by least-squares fitting the experimental values of F for HAwt with the values of F determined by the model. The value $P_0$=1025/μm$^2$ minimized the $\chi^2$ comparing experiment to model, and is also consistent with rough estimates that ~0.5-1% of plasma membrane lipids are PIP2 (1), that half of the PM area is occupied by lipids. This value was then fixed for nonlinear fitting of all HA mutants. The factors by which HA and PIP2 densities are elevated within clusters in comparison to the cell average were determined from experimental values, which also establish their proportionality (See Figure S3). The dissociation constant for PH domain to the PIP2 head group ($K_d$=1 μM) from Murray and McLaughlin (2) was used. Note this is higher than the value (0.21 μM) in Kavran et al., (3) but the same order of magnitude. To convert from molarity to number of molecules per unit area, a factor of $10^3$ is used to account for the 1 nm thick zone within which the lipid head groups are concentrated due to the reduced dimensionality of the membrane (1). That is, within a cubic micrometer of volume ($10^{-15}$ L), at a concentration of 1 μM, the distance between molecules within that volume is calculated, then the number of molecules per unit area is calculated for the same intermolecular distance. All other rate constants containing molarity are converted to number density (number of molecules per square micron), or inverse number density, in a similar way.

*Fitting Measured Spatial Overlap of HA and PH using Model.* A nonlinear least-squares fit of the overlap fraction F, as a function of cell-averaged PAmKate-PH ($K_0$) was performed using three free parameters, an amplitude $A_0$, the reaction equilibrium constant $R_1$ for HA-PIP2 binding, and the total PIP2 concentration $P_0$ within the patch of membrane being simulated. Errors in the measured overlap fraction F were determined from experimental data binned as a function of $K_0$, then parametrized with a linear function, which was then used to calculate errors for each measured F as a function of $K_0$.

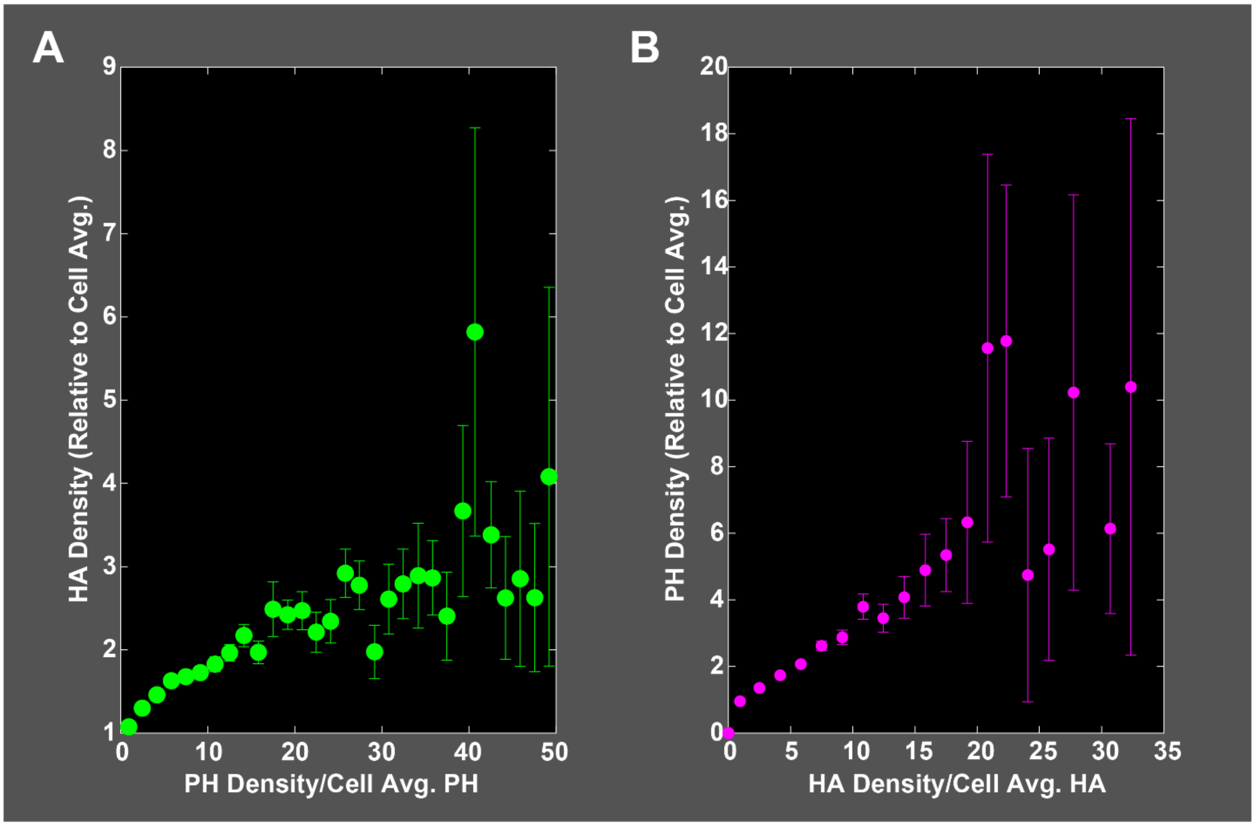


**Figure S3. Nanoscale density of HA and PH are proportional to one another.** Super-resolution microscopy (FPALM) of HA-Dendra2 and PAmKate-PH datasets were quantified to determine the dependence of their density on one another in fixed NIH3T3 cells. The density of localizations (number per square micron) was quantified using a grid of 100 nm square pixels, and then density relative to the cell average determined. (A) The density of HA localizations (green points) as a function of PH density shows an approximately linear trend at PH Densities less than 20 times the cell average, with larger error bars (standard error) at the highest densities where fewer grid pixels contribute to the average. (B) The nanoscale density of PH localizations (magenta points) is approximately linearly dependent on HA density.

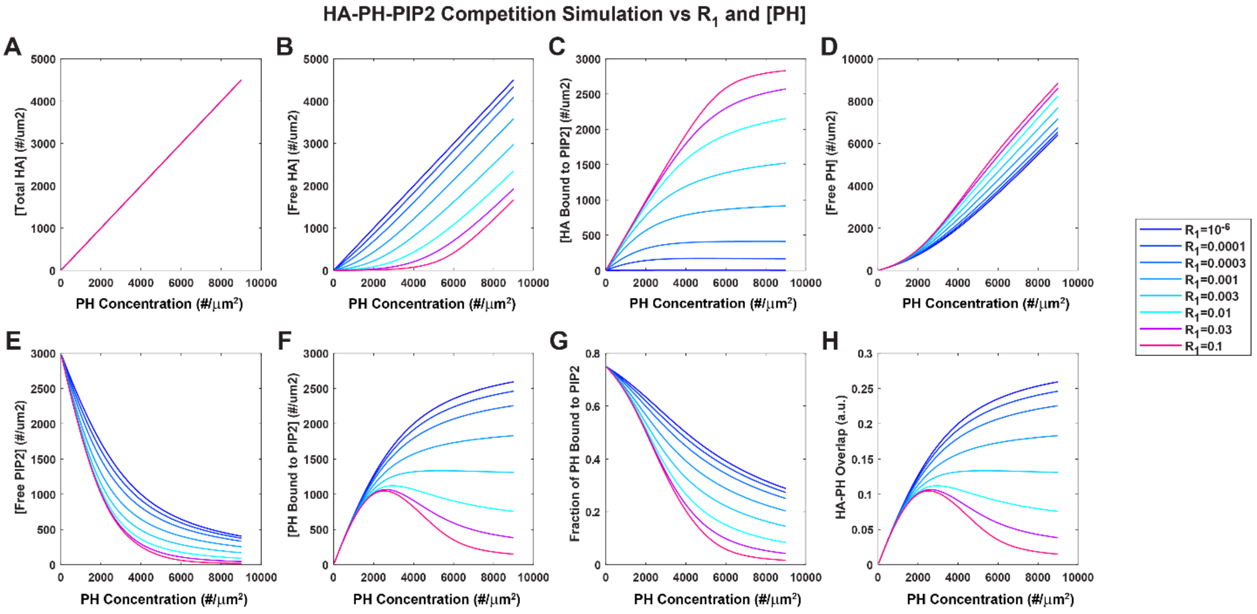


**Figure S4. Predictions of Model of HA-PH-PIP2 Competition.** The analytical solution of the five simultaneous equations (4, 5, 7-9) yields predicted values for A, B, K, P, Q, and HA-PH overlap fraction F as a function of total cell PH concentration and HA-PIP2 binding equilibrium constant $R_1$. Here values are shown for $P_0$ = 3000, $R_2$=0.001, $c_0$=0 and $c_1$=0.5.

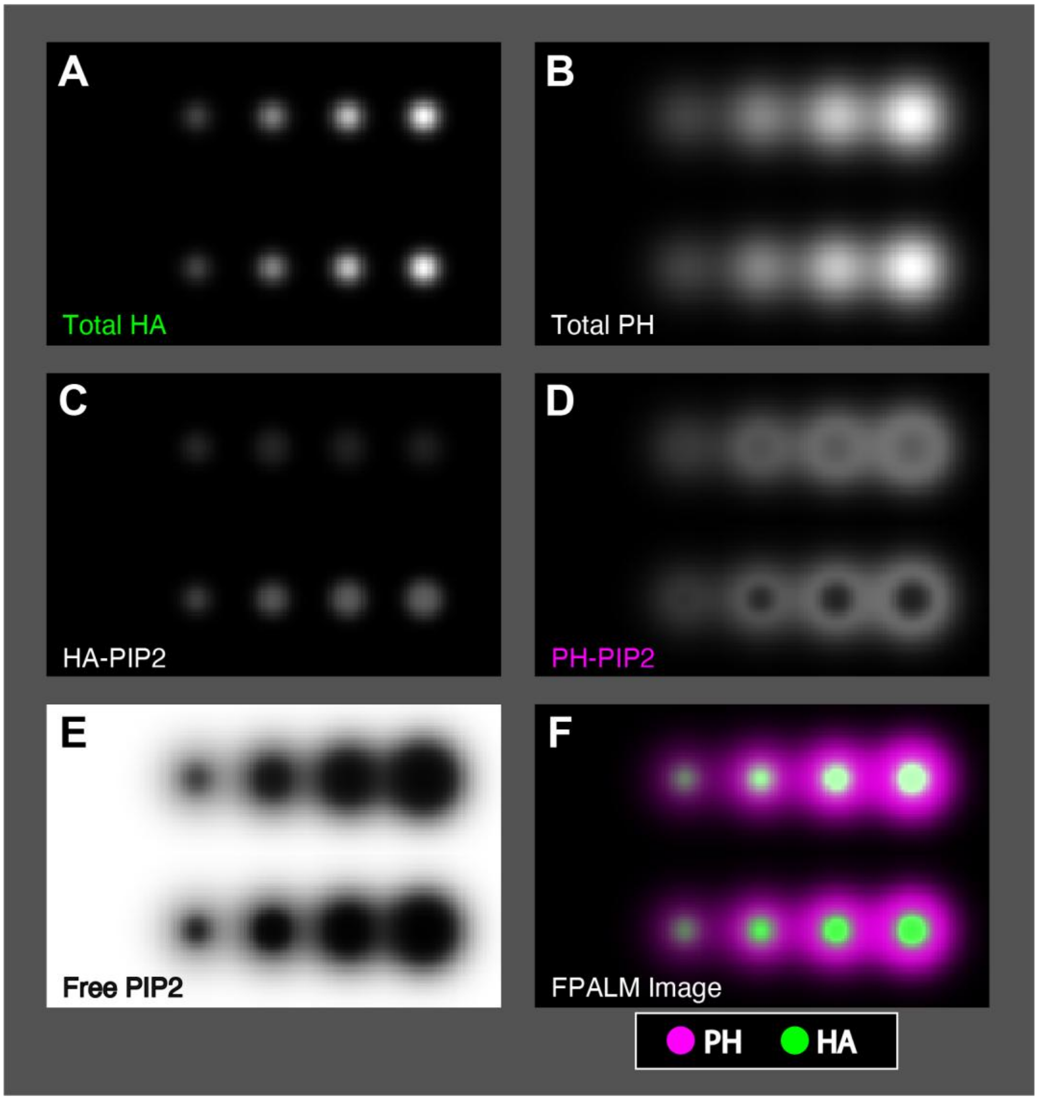


**Figure S5. Simulated FPALM Imaging of HA and PH in HA clusters with weak and strong HA-PIP2 binding**. (A) Small HA clusters with increasing relative density (0, 0.25, 0.5, 0.75, 1.0) from left to right. The upper row of HA clusters has $R_1=R_2/4$, and the lower row of HA clusters has $R_1=R_2 \cdot 4$. (B) Large PH domain clusters surrounding HA clusters with increasing relative density (0, 0.25, 0.5, 0.75, 1.0) from left to right. (C-E) Relative densities predicted by simultaneous equilibrium model for all pixels in the region shown in A and B. (C) HA with PIP2 bound (HA-PIP2) (D) Relative density of PH with PIP2 bound (PH-PIP2) (E) Relative density of free PIP2. (F) Simulated FPALM image of same region shown in A-E, with PH-PIP2 shown in magenta and total HA shown in green. Total PIP2 density for all regions is $2000/\mu m^2$.
